# Dynamic Retuning of Precursor Supply Before and After the Evolution of the Dual Lignin Pathway in Poales

**DOI:** 10.1101/2025.06.05.658145

**Authors:** Jorge El-Azaz, Bethany Moore, Hiroshi A. Maeda

## Abstract

Plants produce diverse and abundant natural products, critical for plant adaptation and human society. While tremendous diversification of plant natural product pathways is well documented, how the upstream primary metabolism evolved in coordination with the lineage-specific pathways remain poorly understood. Here we studied the evolution of aromatic amino acid (AAA) biosynthesis during the emergence of tyrosine-derived lignin pathway uniquely found in grasses and closely related Poales. Phylogeny-guided *in vitro* and *in planta* functional analyses were conducted on two key regulatory enzymes—3-deoxy-D-*arabino*-heptulosonate 7-phosphate synthase (DHS) and arogenate dehydrogenase (TyrA)—catalyzing the first and last steps of AAA and tyrosine biosynthesis, respectively. Deregulated TyrAs were broadly present across Poales, with PACMAD grasses exhibiting stronger deregulation, suggesting that the active tyrosine biosynthesis likely emerged before, but were also selected upon, the evolution of tyrosine-lignin pathway. Conversely, DHS deregulation occurred more recently within BOP grasses after the emergence of tyrosine-lignin pathway, via multiple mutations that acted synergistically in the enzyme’s allosteric regulatory domain. The study highlights how the regulation of upstream primary metabolism is dynamically tuned both before and after the evolution of emerging natural product pathways. These findings underscore the importance of coordinating primary and secondary metabolism for efficient production of natural products in plants.

## INTRODUCTION

Primary metabolites, such as amino acids, are indispensable cellular components that are synthesized through highly conserved pathways. In plants, these primary metabolites also serve as precursor molecules to diverse and abundant lineage-specific natural compounds. Therefore, the primary metabolism pathways must be finely orchestrated with the downstream specialized metabolism, by adjusting different regulatory layers of flux-controlling enzymatic steps. Prior studies have provided comprehensive insight into the importance of transcriptional coordination between enzymes of plant primary and specialized metabolism (van der Fits and Memelink, 2000; Verdonk et al., 2005; Malitsky et al., 2008; Nieuwenhuizen et al., 2015; Ying et al., 2020; Bomal et al., 2013; El-Azaz et al., 2020), which can evolve at a relatively short timescale, such as during crop domestication (Wang et al., 2022a; Maeda and Fernie, 2021). In contrast, we have limited knowledge regarding evolution of enzyme’s post-transcriptional regulation that are needed to increase pathway flux and precursor supply, such as relaxation of feedback inhibition in primary metabolism. This knowledge gap is partly due to the longer timescale of such evolutionary changes (e.g., at family or order level), which makes them difficult to trace back (Maeda and Fernie, 2021). This lack of fundamental understanding limits our ability to engineer plant primary metabolism to enable high and efficient production of natural products and other chemicals in plants.

Phenylpropanoids constitute an abundant and structurally diverse class of plant specialized metabolites that are derived from the aromatic amino acid (AAA) phenylalanine, and uniquely from tyrosine in grass species. In contrast with the high diversity of plant phenylpropanoid metabolism, the upstream AAAs pathway providing the phenylpropanoid precursors, phenylalanine and tyrosine, is largely conserved from bacteria to plants (**Figure 1a**). While dicot plants synthesize lignin and other phenylpropanoids exclusively from phenylalanine via the enzyme phenylalanine ammonia-lyase (PAL) (**Figure 1a**), grasses can utilize both phenylalanine and tyrosine due to the presence of a bifunctional phenylalanine/tyrosine ammonia-lyase (PTAL) enzyme (**Figure 1a**) (Young et al., 1966; Reid et al., 1972; Rösler et al, 1997; Barros et al., 2016; Jun et al., 2018; Barros and Dixon, 2020; Simpson et al., 2021). Grass PTAL emerged from a monofunctional PAL during the evolution of the grass lineage, namely the order Poales (Takeda-Kimura et al., 2024). The ability to synthesize lignin and other phenylpropanoids from tyrosine allows grasses to bypass the cinnamate 4-hydroxylase (C4H) reaction in the early steps of the phenylpropanoid pathway (**Figure 1a**).

**Figure 1.**
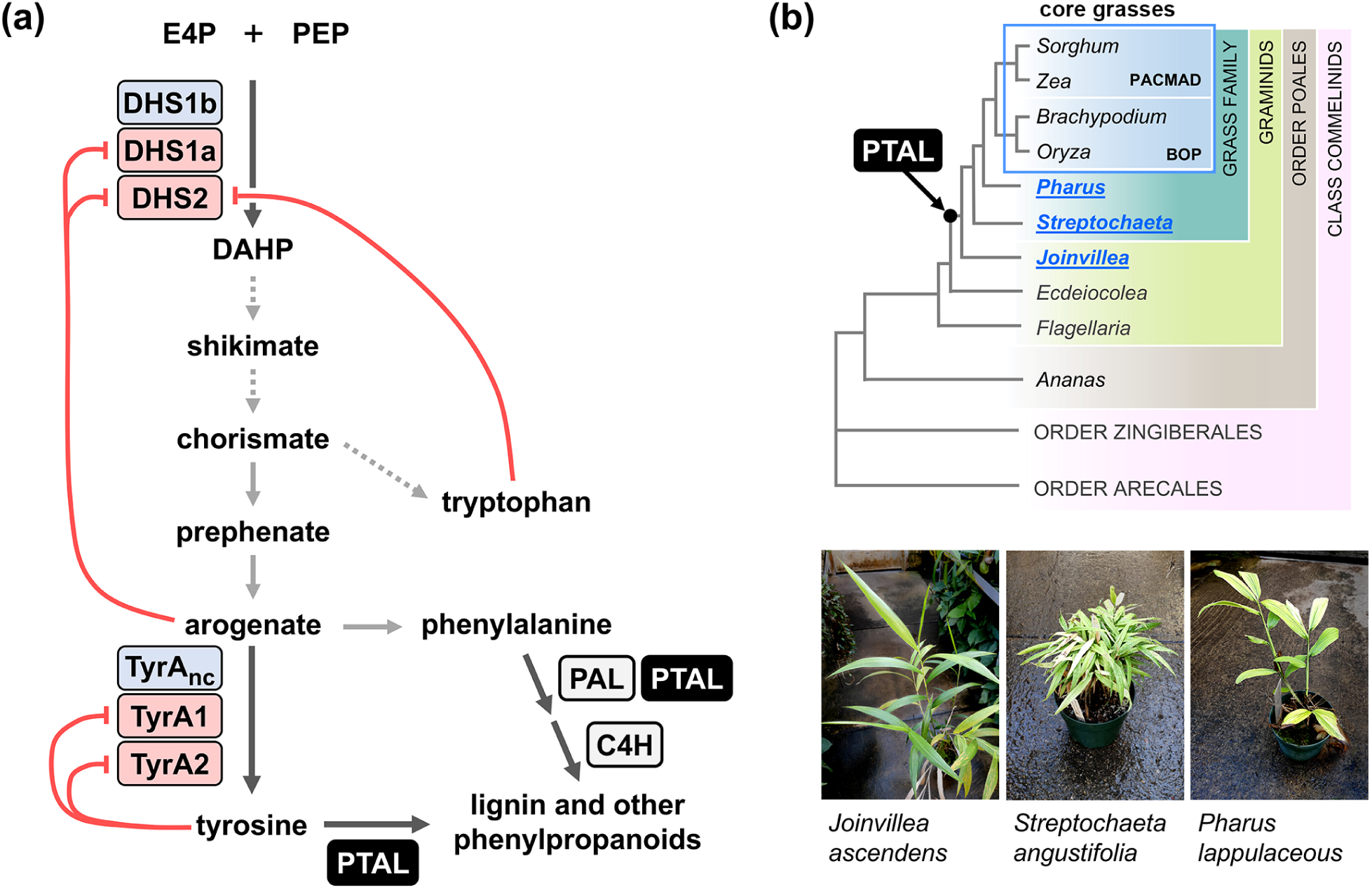
The biosynthesis of aromatic amino acids and the entry steps to phenylpropanoid production in Poales. **(a)** The aromatic amino acid pathway and dual entry phenylpropanoid pathway of grasses. DHS (3-deoxy-D-arabino-heptulosonate 7-phosphate synthase enzyme) and TyrA (arogenate dehydrogenase enzyme) isoforms having different sensitivity to feedback regulation by intermediates and products of the aromatic amino acid pathway are key to modulate the pathway’s flux. Dashed arrows indicate multiple enzymes involved. E4P, erythrose 4- phosphate; PEP, phospho*enol*pyruvate; DAHP, 3-deoxy-D-arabino-heptulosonate 7-phosphate; PAL, phenylalanine ammonia-lyase enzyme; PTAL, bifunctional phenylalanine/tyrosine ammonia-lyase enzyme; C4H, cinnamate 4-hydroxylase. **(b)** Overview of the phylogenetic relationships of PACMAD and BOP grass species compared to other Poales species used in his study (highlighted in blue in the phylogeny), based on McKain et al., 2016 and Givnish et al., 2018. PTAL enzyme likely evolved at some point after the splitting of *Ecdeiocolea sp.* (family Ecdeiocoleaceae) and *Joinvillea sp.* (family Joinvilleaceae) (Takeda-Kimura et al., 2024).

In plants, the feedback regulation of the shikimate and the post-chorismate pathways (**Figure 1a**) controls the production of AAAs, and by extension that of lignin and other phenylpropanoids (Maeda and Dudareva, 2012; Tzin and Galili, 2010). The first enzyme of the shikimate pathway, 3-deoxy-D-*arabino*heptulosonate 7-phosphate synthase (DAHP synthase, or hereafter simply DHS) (**Figure 1a**), is feedback-regulated by AAAs and other derived compounds, effectively controlling how much carbon flows from primary metabolism into the AAAs pathway (Tzin et al., 2012; 2013; Yokoyama et al., 2021; 2022a; 2022b; El-Azaz et al., 2023). In the post-chorismate pathway for tyrosine and phenylalanine production, arogenate dehydrogenase (TyrA) (**Figure 1a**) enzymes are tightly feedback-regulated by tyrosine (Byng et al., 1981; Gaines et al., 1982; Connelly and Conn, 1986; Rippert and Matringe, 2002; Lopez-Nieves et al. 2018; 2022; El-Azaz et al., 2023), which favors the production of phenylalanine—the sole precursor to phenylpropanoids in most plants—over tyrosine. Hence, the cost-efficient grass shortcut to lignin via tyrosine poses an intriguing question regarding the regulation of upstream primary metabolism, as grasses must be able to supply enough tyrosine to feed the tyrosine-derived lignin pathway.

Our previous study found that grass species can support a high rate of tyrosine biosynthesis by means of having partially feedback-insensitive DHS and TyrA enzymatic isoforms, namely BdDHS1b and BdTyrAnc in the model grass *Brachypodium distachyon* (**Figure 1a**) (El-Azaz et al., 2023). It remains unknown, however, when the deregulated BdDHS1b and BdTyrAnc enzymes evolved within the grass lineage, and if their emergence coincides with that of PTAL and the dual phenylalanine/tyrosine lignin pathway. To explore the underlying history of the primary metabolism evolution, here we conducted phylogeny-guided *in vitro* and *in planta* characterization of DHS and TyrA enzymes across diverse species within the order Poales, in which the grass family (Poaceae) belongs (**Figure 1b**). Importantly, this survey includes the grass sister species *Joinvillea ascendens* (family Joinvilleaceae), which has a PTAL enzyme (Takeda-Kimura et al., 2024) and diverged from the grass lineage prior to the *rho* whole genome duplication that affected the grass family (McKain et al., 2016; Givnish et al., 2018). Our findings indicate that feedback-insensitive TyrAnc is an ancient trait of the order Poales that was already present before the emergence of PTAL and the tyrosine-derived lignin pathway. In contrast, deregulated DHS1b emerged more recently within grasses via multiple mutations around the allosteric binding site that synergistically relax DHS feedback regulation. These mutations causing DHS deregulation are not necessarily conserved across grass species, showcasing the diversity of AAAs metabolic regulation across the grass family.

## RESULTS

### Non-canonical TyrA (TyrAnc) is conserved as a single copy gene in Poales

To investigate how tyrosine biosynthesis evolved in Poales, we examined phylogenetic relationships of the TyrA family enzymes. Core grass species generally have three TyrA isoforms (**Figure 1a**), namely TyrA1, TyrA2 and TyrAnc, all of them localized in the plastids (El-Azaz et al., 2023). Whereas grass TyrA1 and TyrA2 share >90% sequence identity between them and are both feedback-inhibited, they are distantly related to TyrAnc, which exhibits relaxed sensitivity to feedback inhibition (El-Azaz et al., 2023; Schenck et al., 2015; 2017; 2018).

To further investigate the phylogenetic relationships of TyrAs in Poales, we reconstructed a detailed phylogenetic tree using TyrA sequences from 32 species from diverse monocot orders, including 18 species from the order Poales (13 grass species in addition to five non-grass Poales), along with *Amborella trichopoda* (**Supplemental Table S2**). The phylogeny included the non-grass graminids *Joinvillea ascendens* (Joinvilleaceae, Takeda-Kimura et al., 2024), *Ecdeiocolea monostachya* (Ecdeiocoleaceae; Takeda-Kimura et al., 2024), and *Flagellaria indica* (Flagellariaceae; McKain et al., 2016), sister lineages to the entire grass family (McKain et al., 2016; Givnish et al., 2018). We also analyzed the TyrAs from the non-core grasses *Streptochaeta angustifolia* (Seetharam et al., 2021) and *Pharus latifolius* (Ma et al., 2021) from the subfamilies Anomochlooideae and Pharoideae, respectively, that are sister to the rest of the grass family (i.e. core grasses; **Figure 1b**) (McKain et al., 2016; Givnish et al., 2018).

This phylogenetic reconstruction confirmed that grasses have three common *TyrA* genes— *TyrA1*, *TyrA2* and *TyrAnc*—with some exceptions in the form of paralog genes derived from more recent duplication events, such as in *Zea mays* (maize) and *Panicum virgatum* (switchgrass, **Figure 2**). In contrast, we found only two full-length *TyrAs* in the genomes of the non-grass Poales and the non-core grasses *Streptochaeta angustifolia* and *Pharus latifolius* (**Figure 2**). These two genes were named as Poales’ *TyrA1* and *TyrAnc*, as they were the outgroup of grass *TyrA1*/*TyrA2* and grass *TyrAnc*, respectively (**Figure 2**). TyrAnc formed an orthogroup already present in *Amborella trichopoda* and conserved in Poales and other monocots (**Figure 2**). Synteny analysis showed that *TyrAnc* is syntenic across graminids, further supporting a common origin (**Supplemental Figure S1**). These findings suggest grass *TyrA1* and *TyrA2* are derived from a duplication event of Poales’ *TyrA1*, which likely took place after the divergence between *Pharus sp*. and the rest of the grass family, whereas TyrAnc appears to have a much more ancient origin.

**Figure 2.**
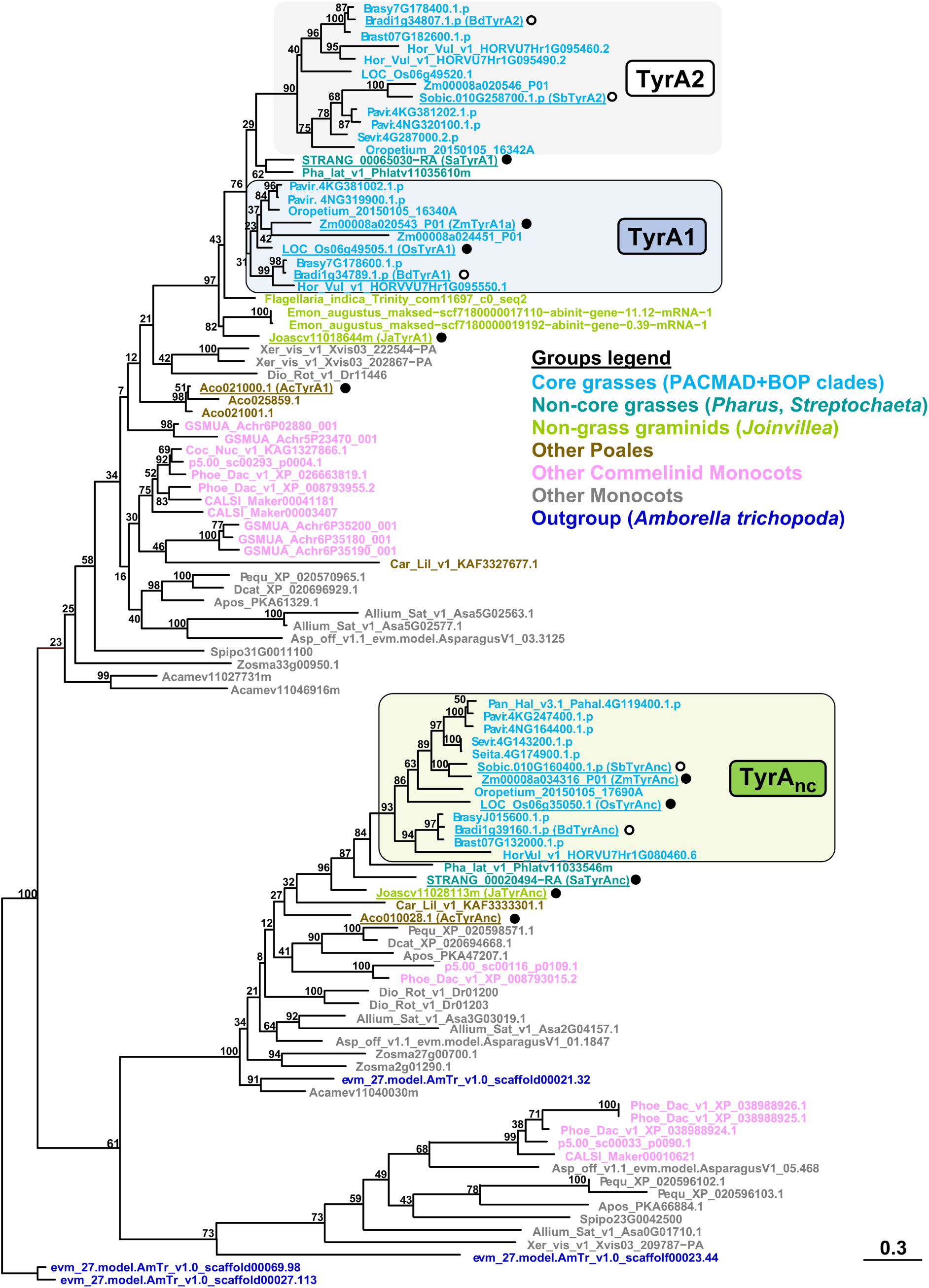
**Phylogenetic reconstruction of TyrA family supports the ancient origin of non- canonical TyrA (TyrAnc) of Poales.** Sequences marked with a solid circle have been characterized in the present study; those marked with a hollow circle were characterized in our previous study (El-Azaz et al., 2023). The phylogeny was inferred from full length protein sequences using maximum likelihood estimation. Bootstrap values were based on 1000 replicates. No outgroup was defined.

### Deregulated TyrAnc enzymes are conserved across Poales

To examine the biochemical properties of TyrAnc orthologs across Poales species, we produced and characterized the recombinant TyrAnc enzymes from *Ananas comosus* (AcTyrAnc), *Joinvillea ascendens* (JaTyrAnc) and *Streptochaeta angustifolia* (SaTyrAnc) (**Figures 1b**, **2**), and two core grass species, each one belonging to one of the two major grass clades: OsTyrAnc, from *Oryza sativa* (rice; BOP clade), and ZmTyrAnc from *Zea mays* (maize; PACMAD clade). As a control for feedback inhibited enzymes, the canonical TyrA1 orthologs from the same five species were also produced and characterized.

These five TyrA1 enzymes shared similar kinetic parameters, such as the Michaelis-Menten constant (*K*m) for the substrate, arogenate and catalytic constant (*k*cat) (**Table 1**; **Supplemental Figure S2**). Similar to what has been reported in TyrAs from other plant species (Byng et al., 1981; Gaines et al., 1982; Connelly et al., 1986; Rippert and Matringe, 2002), all were highly sensitive to feedback inhibition by the reaction product, tyrosine, having *IC*50 values in between 20 to 60 µM when assayed at 500 µM of substrate (**Table 1**; **Figure 3a**). Overall, these kinetic parameters were comparable to those previously reported for TyrA1 in *Brachypodium distachyon* (BdTyrA1) and *Sorghum bicolor* (SbTyrA1) (**Table 1**) (El-Azaz et al., 2023). Therefore, the biochemical properties of TyrA1 enzymes seem to be largely conserved across Poales species, with all TyrA1 analyzed sharing high sensitivity to tyrosine inhibition.

**Figure 3.**
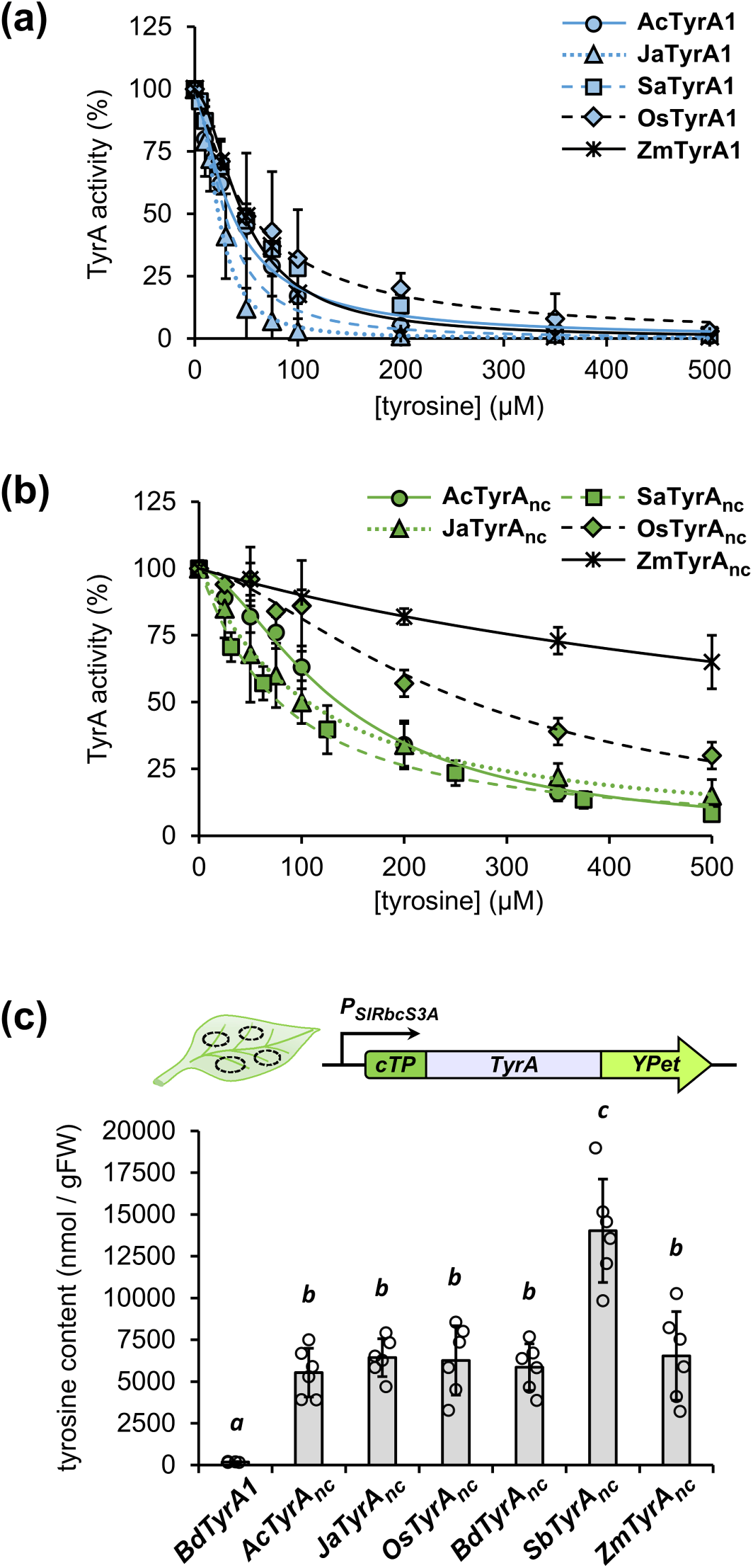
Poales non-canonical TyrA enzymes (TyrAnc) have low sensitivity to feedback inhibition. **(a)** *In vitro* assay for sensitivity to feedback-inhibition (*IC*50 curve) by tyrosine at 0.5 mM of substrate (arogenate) across canonical TyrA1 enzymes from various Poales species: pineapple (AcTyrA1), *Joinvillea ascendens* (JaTyrA1), *Streptochaeta angustifolia* (SaTyrA1); and two core-grass species: rice (OsTyrA1) and maize (ZmTyrA1a). **(b)** *IC*50 curves for tyrosine inhibition in non-canonical TyrA from the same species tested in (a). Datapoints in (a) and (b) represent an average of 3 independent assays. Error bars = *SD*. **(c)** HPLC-FLD determination of tyrosine levels in *Nicotiana benthamiana* leaf tissue expressing *TyrAnc* from diverse Poales species, at 72 hours post-infiltration. *BdTyrA1* was used as control of a grass feedback-inhibited grass TyrA enzyme. Bars represent the average of *n* = 6 biological replicates coming from independent plants. Independent datapoints are represented by hollow circles. Error bars = *SD*. Different letters indicate statistically significant differences (*P* < 0.05) according to Student’s t-test (two-tailed test, equal variance).

In contrast, the kinetic properties of TyrAnc enzymes were diverse across the species analyzed. AcTyrAnc and JaTyrAnc showed high *K*m values towards arogenate (∼1,000 and ∼1,900 µM, respectively, **Table 1**; **Supplemental Figure S2**), whereas SaTyrAnc and the TyrAnc of core- grass species showed 5-10 times lower *K*m values (**Table 1**; **Supplemental Figure S2**). Regarding sensitivity to inhibition by tyrosine, AcTyrAnc, JaTyrAnc, and SaTyrAnc had *IC*50 values of ∼110, ∼130 and ∼85 µM of tyrosine, respectively. However, grass TyrAnc enzymes showed higher *IC*50 values, and were thus more resistant to inhibition by tyrosine: ∼280 µM for OsTyrAnc and ∼850 µM ZmTyrAnc (**Table 1**) (**Figure 3b**). It must be noted that, for each species, *IC*50 of TyrAnc was always at least two-times higher than that of the TyrA1 isoform (**Table 1**). These *in vitro* data show that TyrAnc enzymes of Poales exhibit substantial variations in their kinetic properties, but all sharing relaxed feedback inhibition by tyrosine.

### **Expression of TyrAnc from PACMAD grasses yield the highest tyrosine levels in** *Nicotiana benthamiana*

Next, we investigated whether the observed differences in the biochemical properties of TyrAnc from different Poales impact tyrosine production *in planta*. To this end, we performed Agrobacterium-mediated transient expression of *AcTyrAnc*, *JaTyrAnc*, *SaTyrAnc, BdTyrAnc*, *SbTyrAnc* and *ZmTyrAnc* in the leaves of *Nicotiana benthamiana* expressed under control of the CaMV 35S promoter (**Supplemental Figure S3**). Using these constructs, tyrosine content in the plant samples did not correlate with the *in vitro* sensitivity to tyrosine inhibition of the TyrAnc enzymes (**Supplemental Figure S4**). Determination of TyrA protein levels by quantitative anti-FLAG tag immunoblot revealed high variability, up to 100-times, in their abundance between different TyrAnc tested (**Supplemental Figure S4**), likely precluding quantitative *in planta* comparison of different TyrAs. We speculated that this limitation could be overcome by 1) reducing the intensity of the overexpression by using a weaker promoter and 2) stabilizing the enzymes by fusing them to a GFP-family tag, since we were able to produce BdTyrAnc-EGFP at high levels in Arabidopsis protoplasts previously (El-Azaz et al., 2023).

We therefore expressed *AcTyrAnc*, *JaTyrAnc*, *OsTyrAnc*, *BdTyrAnc*, *SbTyrAnc* and *ZmTyrAnc* fused to *YPet* (a YFP variant) in their C-terminus and under control of the RuBisCO small subunit 3A promoter from tomato (*PSlRbcS3A*; **Supplemental Figure S3**), which provides approximately 15 to 20% of the expression level of *CaMV* 35S promoter (Engler et al., 2014). *BdTyrA1*-*YPet* was also expressed as a control for a feedback-inhibited grass enzyme. Laser scanning confocal microscopy of the infiltrated *Nicotiana benthamiana* leaves confirmed plastidial localization for all fusion proteins (**Supplemental Figure S5**). Quantification of TyrA-YPet fusion proteins by immunoblot at 72 hours post-infiltration showed a drastic reduction in the variability among TyrA orthologs compared to the previous experiment, ranging now from ∼110 to ∼140 µg per gFW for all TyrAnc proteins, except ZmTyrAnc, which remained low abundance at ∼4.5 µg/gFW (**Supplemental Figure S6**).

Tyrosine content, determined by high performance liquid chromatography (HPLC), was similar in *AcTyrAnc*, *JaTyrAnc, OsTyrAnc* and *BdTyrAnc* infiltrated leaves, around 4,000-6,000 nmol of tyrosine per gFW, ∼50-times higher than the *BdTyrA1* control (**Figure 3c**). Leaves expressing *SbTyrAnc* rounded 12,000 nmol/gFW of tyrosine, significantly higher than tyrosine levels in leaves expressing *TyrAnc* enzymes of Poales and BOP grasses (**Figure 3c**). Despite the low abundance of ZmTyrAnc protein (∼20-times lower than the other TyrAnc tested), tyrosine levels upon expressing *ZmTyrAnc* were equivalent to *AcTyrAnc*, *JaTyrAnc*, *OsTyrAnc* and *BdTyrAnc* infiltrated leaves, which is consistent with the *in vitro* data showing strong de-regulation of ZmTyrAnc (i.e., high *IC*50 and *K*i; **Table 1**). Combined with biochemical evidence, these findings suggest that partially deregulated TyrAnc enzymes are conserved across Poales, with the strongest deregulation observed in orthologs from PACMAD grasses.

### **Poales’ single copy** *DHS1* **duplicated into** *DHS1a* **and** *DHS1b* **within the grass family**

To track the evolutionary origin of deregulated DHS1b of grasses, we performed a phylogenetic reconstruction of the *DHS* genes from Poales and other monocots, using the same set of species as in the TyrA phylogeny (**Supplemental Table 2**). The resulting DHS phylogeny supported the existence of four groups of DHS sequences in core grass species (*DHS1a*, *DHS1b*, *DHS2* and *DHSnc*; El-Azaz et al., 2023), with grass *DHS1a* and *DHS1b* being closely related between them (**Figure 4**).

**Figure 4.**
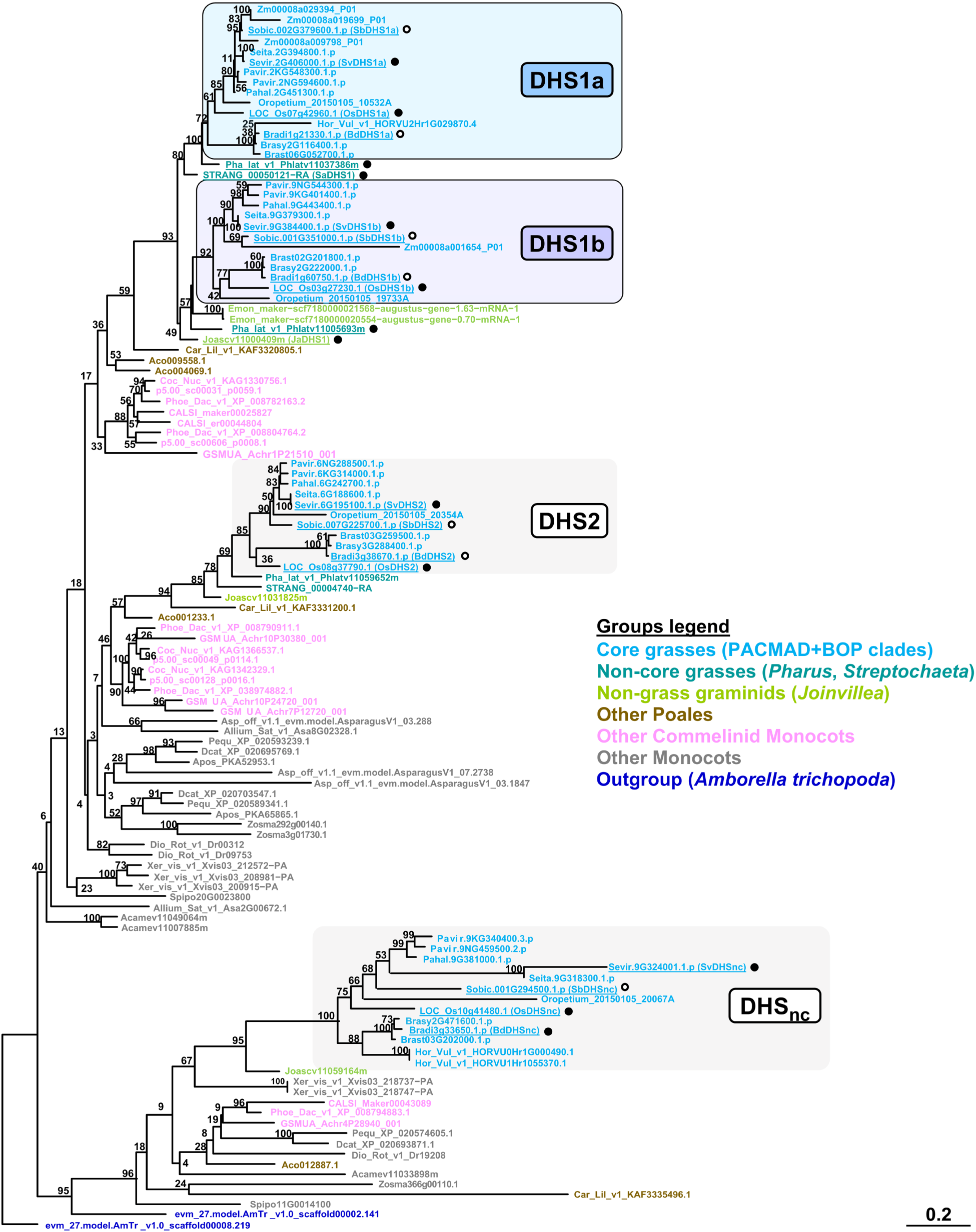
Phylogenetic reconstruction of DHS family supports the recent emergence of deregulated DHS1b enzyme of grasses. Sequences marked with a solid circle have been characterized in the present study; those marked with a hollow circle were characterized in our previous study (El-Azaz et al., 2023). The phylogeny was inferred from full length protein sequences using maximum likelihood estimation. Bootstrap values were based on 1000 replicates. No outgroup was defined.

Contrary to the four *DHS* genes found across core grasses, the graminid *Joinvillea ascendens* and the non-core grass *Streptochaeta angustifolia* had three *DHS* genes. These species had only one DHS copy in the *DHS1a*/*DHS1b* clade, corresponding with the gene models *Joascv11000409m* in *Joinvillea ascendens* and *STRANG_00050121−RA* in *Streptochaeta angustifolia* (**Figure 4)**, thus named *JaDHS1* and *SaDHS1*, respectively. In contrast, the genome of the non-core grass *Pharus latifolius*, sister to the rest of the grass family (i.e., core-grasses), harbored two *DHS1a* and *DHS1b* orthologs (**Figure 4**). These observations suggest that *DHS1a* and *DHS1b* emerged from the duplication of a Poales’ single copy *DHS1* before the divergence between *Pharus* sp. and core grasses.

### Poales’ DHS1 and non-core grass DHS1b enzymes have low *in planta* activity

To examine the feedback regulation of the DHS enzyme encoded by the single copy *DHS1* of Poales, we determined the sensitivity of JaDHS1 and SaDHS1 to effector-mediated inhibition. *JaDHS1* and *SaDHS1* were cloned into the pET28a vector, and the activity of the purified recombinant proteins determined at 0.5 mM of each AAA or arogenate, which inhibit BdDHS1a but not BdDHS1b *in vitro* (El-Azaz, 2023). These experiments revealed that JaDHS1 and SaDHS1 were inhibited by arogenate (**Figure 5a**), suggesting that the Poales’ DHS1 are feedback-inhibited enzymes.

**Figure 5.**
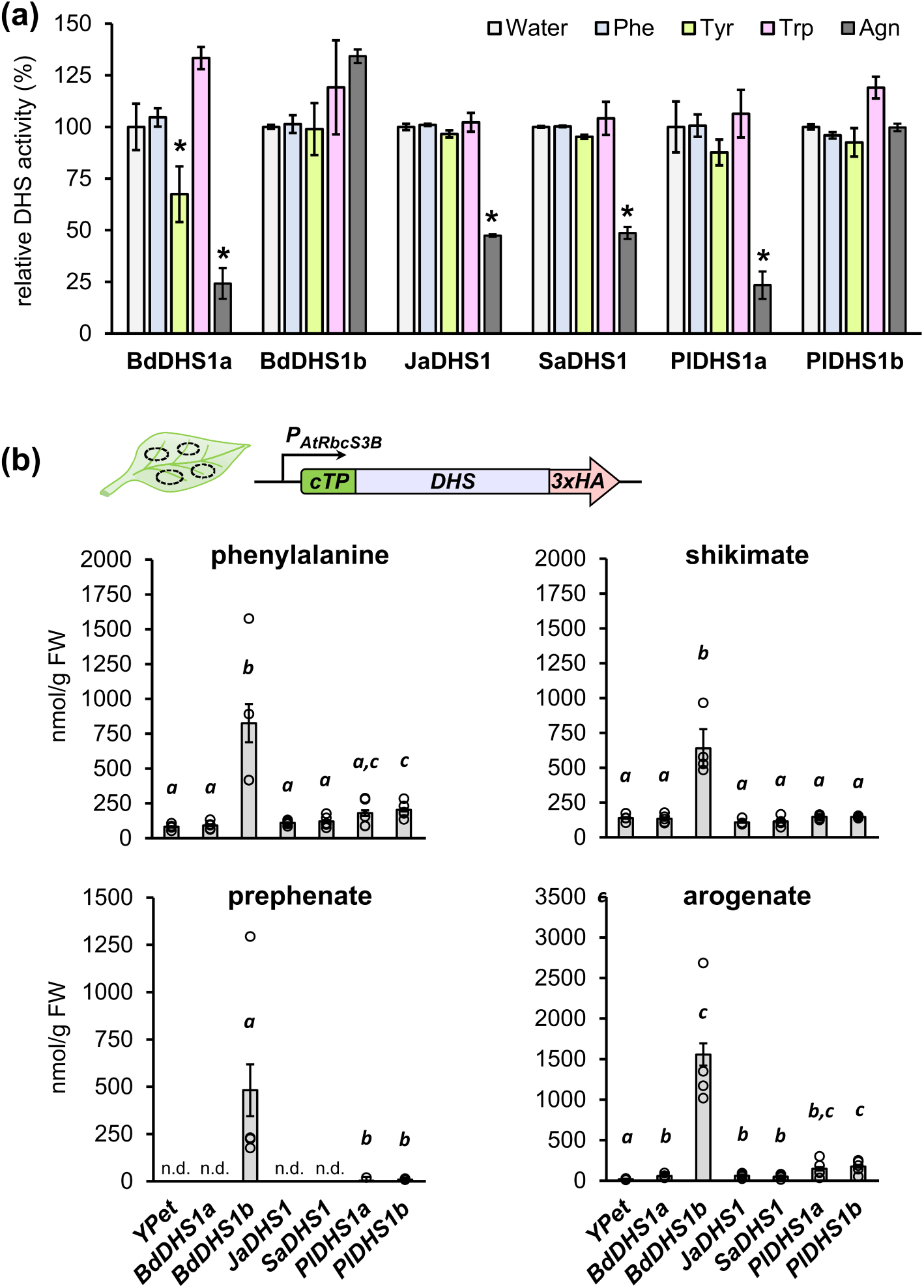
Deregulated DHS1b emerged after the split between Pharus sp. and the rest of the grass family. **(a)** DHS phylogeny in Commelinid monocots, including grasses. Species color code as in figure 2a **(b)** Recombinant DHS1a/1b orthologs from *Brachypodium distachyon* (BdDHS1a and BdDHS1b), *Joinvillea ascendens* (JaDHS1), *Streptochaeta angustifolia* (SaDHS1) and *Pharus lappulaceous* (PlDHS1a and PlDHS1b) were assayed side by side at 0.5 mM of phenylalanine (Phe), tyrosine (Tyr), tryptophan (Trp) or arogenate (Agn) as effectors. BdDHS1a and BdDHS1b were used as controls. Water was used as a no-effector control to determine 100% of the relative activity of each enzyme. Asterisks (*) indicate significantly lower DHS activity compared to the corresponding water control, according to Student’s t-test (two-tails test, equal variance). **(c)** Levels of phenylalanine, and its precursors shikimate, prephenate and arogenate, as determined by LC-MS in the leaves of *Nicotiana benthamiana* at 72 hours post-infiltration of constructs expressing *JaDHS1*, *SaDHS1*, *PlDHS1a* and *PlDHS1b*. *BdDHS1a* and *BdDHS1b* were included as controls. Note that different scales have been used for the different compounds. Data presented as the average of *n* = 5 samples coming from independent plants, except *n* = 4 for the *YPet* negative control and *BdDHS1b*. Error bars = *SE*. Different letters indicate statistically significant differences (P < 0.05) according to Student’s t-test (two-tailed test, equal variance). n.d. = not detected.

To narrow down the evolutionary timing of the emergence of functionally de-regulated DHS1b, we characterized the DHS1a and DHS1b enzymes from the non-core grass *Pharus sp.*, the first graminid species having both DHS1a and DHS1b orthologs (**Figure 4**). As we could not have access to *Pharus latifolius* plant tissue (for which the genome sequence is available), we cloned these two genes from the closely related species *Pharus lappulaceous,* namely PlDHS1a and *PlDHS1b* (GeneBank identifiers PV055691 and PV055692, respectively), which shared 99.8% and 98.6% protein sequence identity, respectively, with the *Pharus latifolius* orthologs (excluding the plastid transit peptide). The *in vitro* characterization of these two proteins showed that PlDHS1a is inhibited by arogenate, like grasses DHS1a, but PlDHS1b was not inhibited by AAAs or arogenate (**Figure 5a**). These results indicate that DHS1b from *Pharus lappulaceous* is a deregulated enzyme.

To contrast these *in vitro* findings, *JaDHS1*, *SaDHS1*, *PlDHS1a* and *PlDHS1b* were subcloned into the Golden Gate plant expression vector pICH47822 under control of the promoter of Arabidopsis RuBisCO small subunit 3B (*PAtRbcS3B*) (Engler et al., 2014) (**Supplemental Figure S3**). The 3x Human influenza hemagglutinin (HA) tag was fused in frame to the C-terminus of the *DHS* genes to confirm the production of the heterologous protein. Quantification of the levels of phenylalanine, and its precursors shikimate, prephenate and arogenate by Liquid chromatography–mass spectrometry (LC-MS) at around 72-hours post-infiltration showed that *JaDHS1* and *SaDHS1* expression had little or no impact over the levels of these metabolites compared to the *YPet* negative control (**Figure 5b**). This stays in sharp contrast to the strong effect of *BdDHS1b* expression (**Figure 5b**). Quantification of JaDHS1 and SaDHS1 HA fusion proteins by immunoblot showed that these two proteins were accumulated at 3- to 5-times higher levels than BdDHS1b (**Supplemental Figure S7**). Therefore, these data together support that, unlike BdDHS1b, JaDHS1 and SaDHS1 have very low activity *in planta*.

Unexpectedly, *PlDHS1a* and *PlDHS1b,* which showed different feedback regulation *in vitro* (**Figure 5a**), both had a similar effect *in planta*, only causing relatively moderate increases in phenylalanine (by two-times) and arogenate (by seven-times) levels compared to the *YPet* negative control (**Figure 5b**). In sharp contrast, the feedback-insensitive *BdDHS1b* increased the levels of phenylalanine, shikimate and arogenate by 10-, 5- and 75-times, respectively, compared to *YPet*, and drastically increased prephenate content (which was otherwise below detection levels; **Figure 5b**). While PlDHS1b is not sensitive to arogenate or tryptophan inhibition *in vitro* (**Figure 5a**), this enzyme is still seemingly much less active *in planta* than BdDHS1b.

### **DHS1b enzymes highly active** *in planta* **emerged within BOP grasses**

To comprehensively explore the distribution of *in planta* deregulated DHSs from grasses, the *in planta* transient expression approach was extended to all four members of the *DHS* gene family (*DHS1a*, *DHS1b*, *DHS2*, *DHSnc*; **Figure 4**) from two BOP clade (*Brachypodium distachyon* and *Oryza sativa*) and two PACMAD clade (*Sorghum bicolor* and *Setaria viridis*) core grass species. All 16 *DHS* genes were again expressed as 3xHA C-terminus fusion proteins under control of *PAtRbcS3B*.

LC-MS analysis of *Nicotiana benthamiana* leaf tissue expressing these *DHS* genes side-by- side, plus *YPet* as negative control, revealed that only *BdDHS1b* and *OsDHS1b* induced strong increases in the levels of shikimate, prephenate and arogenate (**Figure 6**). *BdDHS1b* expression increased shikimate content by around 10-times, prephenate by 300-times, and arogenate by 100- times (**Figure 6**), whilst *OsDHS1b* increased shikimate, prephenate and arogenate levels by approximately 5-, 100- and 40-times, respectively (**Figure 6**), all relative to *YPet* samples. On the other hand, *SvDHS1b* increased shikimate, prephenate and arogenate content by only 2-, 4- and 10-times compared to *YPet* (**Figure 6**), with even lesser effects being induced by *SbDHS1b* expression (**Figure 6**). Quantification of the DHS-HA tagged proteins by immunoblotting showed that, despite having a markedly different impact on metabolite levels, BdDHS1b and SvDHS1b expressed at similar levels *in planta*, at around 30 µg/gFW (**Supplemental Figure S8**). None of the other *DHSs* caused drastic increases in the levels of the target metabolites (**Figure 6**). Therefore, within the grass lineage, highly active DHSs *in planta* appear to be specific to DHS1b orthologs from BOP clade species. This finding indicates that the deregulated DHS enzymes evolved relatively recently as compared to the earlier emergence of deregulated TyrAnc enzymes.

**Figure 6.**
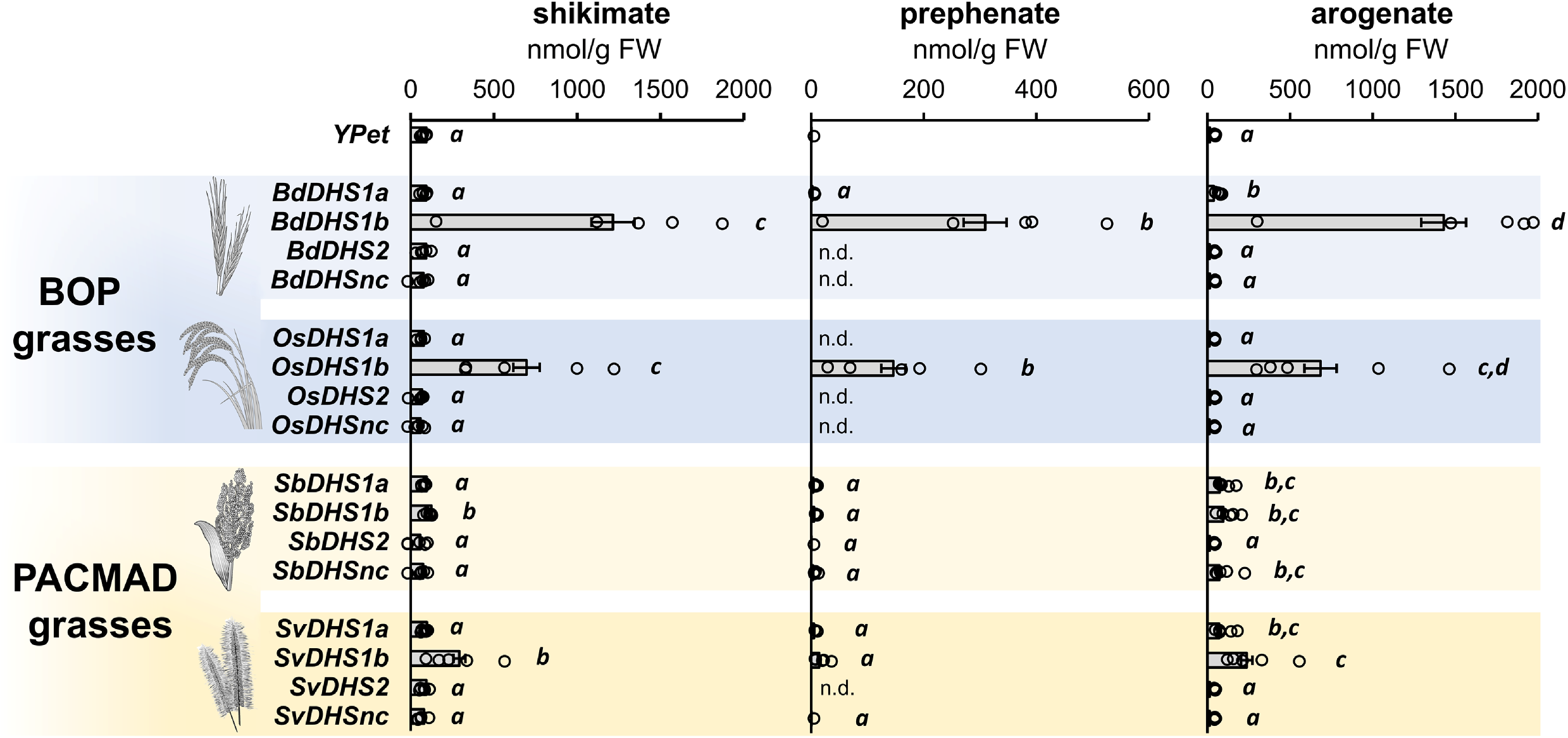
Deregulated DHS1b is limited to BOP grass species. The levels of aromatic amino acid pathway intermediates shikimate, prephenate and arogenate, determined by LC-MS in the leaves of *Nicotiana benthamiana* at three days post-infiltration. The plants were infiltrated with independent constructs expressing each one of the four grass *DHS* genes from two BOP grass species, *Brachypodium distachyon* (*Bd*) and *Oryza sativa* (*Os*), and two PACMAD grasses, *Setaria viridis* (*Sv*) and *Sorghum bicolor* (*Sb*). Note that the scale used to represent prephenate levels is different from that of shikimate and arogenate. Data presented as the average of *n* = 5 samples coming from independent plants. Letters indicate significant statistical groups (*P* < 0.05) according to Student’s *t*-test (two-tailed test, equal variance) . n.d. = not detected.

### Multiple mutations around the allosteric binding site synergistically increase DHS activity *in planta*

A previous genetic screening conducted in our laboratory identified a set of point mutations that cause deregulation of Arabidopsis DHS enzymes (Yokoyama et al., 2022a). Many of these point mutations concentrated in a particular region of the plant DHS enzymes, a helix-loop-helix motif—namely *sota* domain—adjacent to the allosteric regulatory site of plant DHSs (Yokoyama et al., 2022a). To understand what changes in the sequence of Brachypodium and rice DHS1b make these enzymes highly active *in planta*, we first analyzed if any of the Arabidopsis *sota* mutations may be present in BdDHS1b or OsDHS1b. A Thr211Met substitution was found in the *sota* domain of BdDHS1b, but was absent in OsDHS1b, making it uncertain whether Thr211Met alone could be responsible for enhanced DHS activity (**Supplemental Figure S9**).

To identify which other residues contribute to the high activity of BdDHS1b and OsDHS1b *in planta*, we performed a site-directed mutagenesis analysis on PlDHS1b, as it shares near 90% sequence identity with BdDHS1b but shows low *in planta* activity (**Figure 5c**). Using multiple sequence alignments, we selected a set of 17 candidate residues on PlDHS1b that were highly conserved in feedback inhibited enzymes from Poales and grasses, but differed in BdDHS1b and OsDHS1b (**Supplemental Figure S10**). These residues were grouped into three sections or blocks of the protein primary sequence, named MUT1, MUT2 and MUT3 (**Figure 7a**), and were replaced by the corresponding residues from BdDHS1b. Then, these mutated gene blocks were synthesized and assembled to introduce all 17 residue substitutions simultaneously, or combined one by one with the equivalent non-mutated (i.e., wild type) blocks (**Figure 7a**) (**Supplemental Figure S10**).

**Figure 7.**
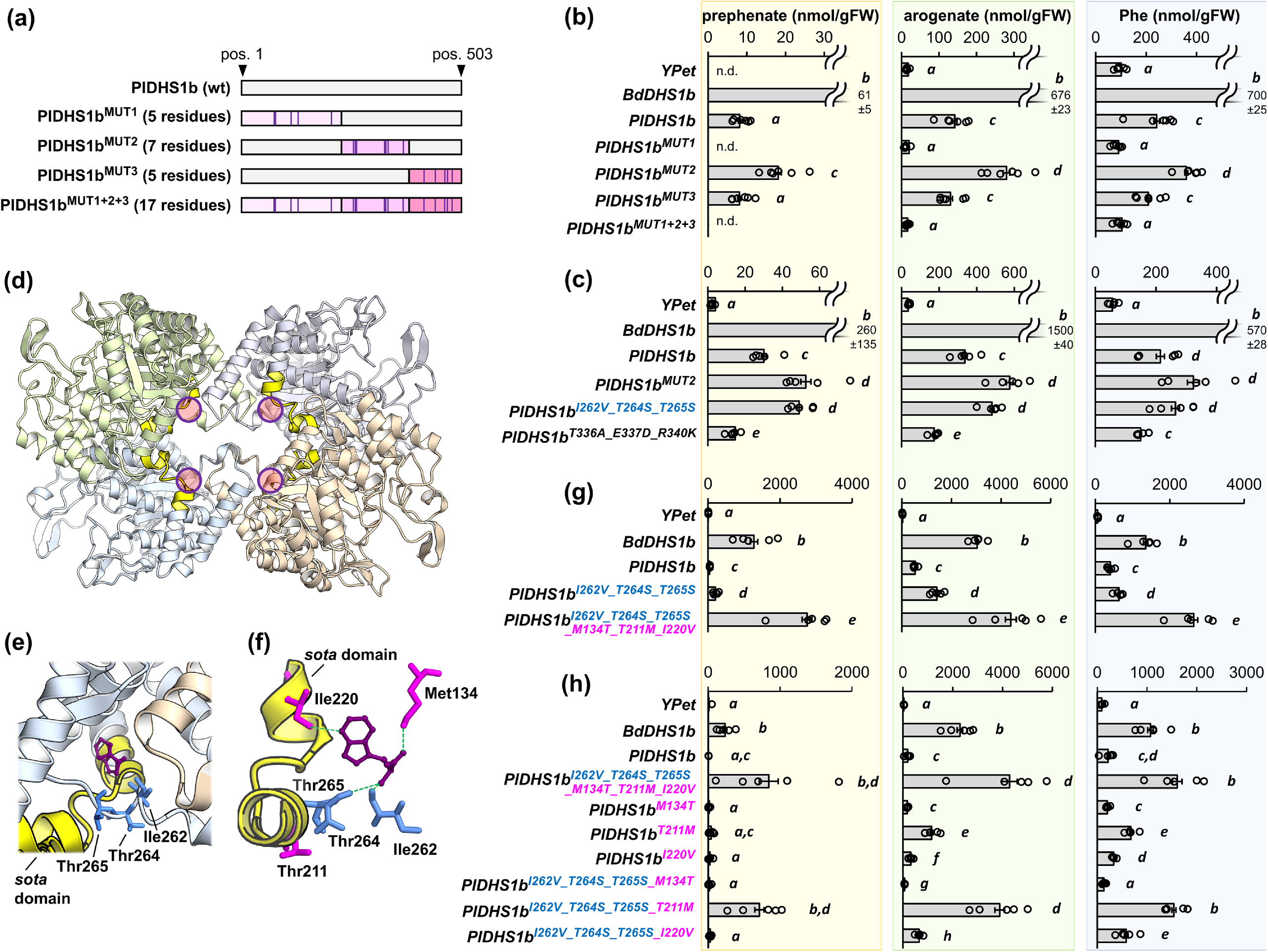
Phylogeny- and structure-guided site-directed mutagenesis on PlDHS1b identifies residues deregulating DHS activity *in planta*. **(a)** Dissection of *Pharus lappulaceous* DHS1b protein (PlDHS1b) into three blocks of mutations, namely MUT1, MUT2 and MUT3. The residues were substituted for the equivalent residue in BdDHS1b sequence, and are marked with purple lines on the models. Residue numeration corresponds to PlDHS1b sequence without the predicted transit peptide. **(b)** *In planta* levels of prephenate, arogenate and phenylalanine upon expression of *PlDHS1b* mutant proteins *PlDHS1b^MUT1^*, *PlDHS1b^MUT2^*, *PlDHS1b^MUT3^* and the triple mutant *PlDHS1b^MUT1+2+3^*, determined at 3-days post-infiltration, compared to wild type PlDHS1b and deregulated BdDHS1b. **(c)** Metabolite levels after transient expression of two lower-order mutant versions of *PlDHS1b^MUT2^*, *PlDHS1b^I262V_T264S_T265S^* and *PlDHS1b^T336A_E337D_R340K^*, units like in (b). **(d)** *In silico* model of PlDHS1b wild type protein based on the tetrameric crystal structure of the *Mycobacterium tuberculosis* DHS (MtDAH7PS) co-crystalized with its effector molecules, phenylalanine and tryptophan. Red circles indicate the analogous location of the allosteric binding site for tryptophan, adjacent to the *sota* regulatory domain previously identified in Arabidopsis DHSs (helix-loop-helix marked in bright yellow). **(e)** and **(f)**, the modelled PlDHS1b tryptophan binding pocket. The putatively bound tryptophan molecule is represented in dark purple, the neighboring residues differing between PlDHS1b and BdDHS1b are highlighted in either blue or magenta (residues and numbering corresponds to PlDHS1b wild type protein without the predicted transit peptide). **(g)** Metabolite levels after transient expression of *PlDHS1b* sextuple mutant *PlDHS1b^M134T_T211M_I220V_I262V_T264S_T265S^*, compared to other PlDHS1b mutant versions and BdDHS1b. **(h)** Dissection of the individual contribution of Met134Thr, Thr211Met and Ile220Val to PlDHS1b deregulation. Data in (b), (c), (g) and (h) presented as the average of *n* = 5 samples coming from independent plants. Note that different scales have been used for the different compounds. Error bars = *SE*. Different letters indicate statistically significant differences (*P* < 0.05) according to Student’s *t*-test (two-tailed test, equal variance). n.d. = not detected.

The resulting four *PlDHS1b* mutated versions, namely *PlDHS1b^MUT1^*, *PlDHS1b^MUT2^*, *PlDHS1b^MUT3^*, and *PlDHS1b^MUT1+2+3^* were then transiently expressed into *Nicotiana benthamiana* following the exact same approach as above. LC-MS analysis of tissue expressing *PlDHS1b^MUT3^* showed no significant effect compared to *PlDHS1b* expression (**Figure 7b**), whereas the expression of *PlDHS1b^MUT1^* and the triple mutant *PlDHS1b^MUT1+2+3^* equaled to the expression of the *YPet* negative control (i.e., markedly lower than *PlDHS1b* wild type), suggesting that MUT1 abolishes enzyme’s activity (**Figure 7b**). On the other hand, only the tissue expressing *PlDHS1b^MUT2^* showed elevated levels of prephenate, arogenate and phenylalanine, by around 2-

, 2- and 1.5-times, respectively, compared to the *PlDHS1b* wild type control (**Figure 7b**). Combining blocks MUT2 and MUT3 into *PlDHS1b^MUT2+3^* mutant or mutating a set of selected 9 residues within the MUT2+3 region did not result in any significant increase in metabolite content compared to *PlDHS1b* (**Supplemental Figure S10 and S11**). Hence, we concentrated our efforts in dissecting the mutations within MUT2.

The MUT2 block comprises seven mutated residues: Ile262Val, Thr264Ser, Thr265Ser, Thr336Ala, Glu337Asp, Arg340Lys, and Arg377His (**Supplemental Figure S10**). To identify which mutation(s) within this region are responsible for the increase in DHS activity, we dissected MUT2 by generating two new mutant versions of PlDHS1b, each one having a mutated triad of closely located residues: PlDHS1b^I262V_T264S_T265S^ and PlDHS1b^T336A_E337D_R340K^. Transient expression of these two mutants revealed that the three mutations—Ile262Val, Thr264Ser, and Thr265Ser— were sufficient to achieve the effect of PlDHS1b^MUT2^ (**Figure 7c**). In contrast, *PlDHS1b^T336A_E337D_R340K^* infiltrated tissue exhibited reduced metabolite levels compared to wild type *PlDHS1b* (**Figure 7c**). Importantly, however, *PlDHS1b^MUT2^* and *PlDHS1b^I262V_T264S_T265S^* were still clearly less active than *BdDHS1b* (**Figures 5, 6, 7a,b**), indicating that Ile262Val, Thr264Ser, and Thr265Ser mutations increase DHS activity *in planta* but are clearly insufficient to reach BdDHS1b and OsDHS1b activity levels.

*In silico* modeling of PlDHS1b protein (**Figure 7d**) using the crystal structure of the tetrameric DHS enzyme from *Mycobacterium tuberculosis* (Protein Data Bank structure 3RZI, named MtDAH7P; Jiao et al., 2012) bound to its allosteric effector molecules, phenylalanine and tryptophan, suggested that residues Ile262, Thr264, and Thr265 form a triad on one side of the allosteric binding site responsible for tryptophan-mediated regulation in MtDAH7P (**Figures 7e,f**) (Jiao et al., 2012). Based on the model, these three residues would be oriented towards the amino group of the allosteric tryptophan molecule (**Figure 7f**). Furthermore, these residues—and particularly Thr265—are structurally adjacent to the helix-loop-helix structure (i.e., the *sota* domain) previously identified in Arabidopsis DHS mutants (Yokoyama et al., 2022a).

To identify additional mutations that may further deregulate PlDHS1b to the level of BdDHS1b, we concentrated our attention on the region surrounding the putative tryptophan allosteric binding site, including the *sota* domain. A careful comparison of this region between PlDHS1b and BdDHS1b pointed to three residues that differ between these two enzymes: Met134, Thr211, and Ile220 in PlDHS1b (**Figure 7f**), corresponding to Thr, Met and Val, respectively, in BdDHS1b (**Supplemental Figure S9**). Whereas Ile220Val was not tested above, Met134Thr and Thr211Met were already mutated within PlDHS1b^MUT1^ (**Supplemental Figure S10**), a protein seemingly devoid of *in planta* activity (**Figure 7b**). Noticeably, residues at positions 211 and 220 were located within the *sota* domain (**Figure 7f**), whilst residue at position 134 was located on the opposite side of the allosteric pocket, possibly oriented towards the effector molecule (**Figure 7f**).

We then introduced these three additional mutations—Met134Thr, Thr211Met and Ile220Val—into the partially de-regulated PlDHS1b^I262V_T264S_T265S^ background, generating the sextuple mutant PlDHS1b^M134T_T211M_I220V_I262V_T264S_T265S^. Expression of this sextuple mutant resulted in drastic increases in metabolite levels: roughly 70-times for prephenate, 9-times for arogenate, and 7-times for phenylalanine, compared to the wild type PlDHS1b (**Figure 7g**). Quantitative immunoblot confirmed equivalent accumulation levels for the PlDHS1b wild type and PlDHS1b^M134T_T211M_I220V_I262V_T264S_T265S^ HA fusion proteins, at around 24 and 29 µg per gFW, respectively (**Supplemental Figure S12**). Metabolite levels ensuing expression of PlDHS1b^M134T_T211M_I220V_I262V_T264S_T265S^ were even higher, around twice, than those observed in leaves expressing BdDHS1b side-by-side (**Figure 7g**).

We further dissected the effect of Met134Thr, Thr211Met and Ile220Val mutations individually by introducing them into both PlDHS1b wild type and PlDHS1b^I262V_T264S_T265S^ (**Figure 7h**). The Met134Thr substitution did not enhance activity of PlDHS1b or PlDHS1b^I262V_T264S_T265S^ (**Figure 7h**). Conversely, compared to PlDHS1b, PlDHS1b^T211M^ increased arogenate and phenylalanine levels by 6- and 3-times, respectively, whilst PlDHS1b^I220V^ increased arogenate levels only by 1.5- times (**Figure 7h**). Notably, a marked additive effect was observed when these mutations were introduced into the PlDHS1b^I262V_T264S_T265S^ background; the quadruple mutant PlDHS1b^I262V_T264S_T265S_T211M^ was particularly effective and became more active than BdDHS1b (**Figure 7h**). Therefore, multiple substitutions around the allosteric binding site synergistically contribute to modulate DHS activity *in planta*, with the combination of Ile262, Thr264, Thr265 and Thr211Met substitutions generating a highly active DHS enzyme *in planta*.

## DISCUSSION

Diverse studies have shown that plants coordinate the expression of primary and specialized metabolism genes in specific tissues and conditions to efficiently synthesize natural products (van der Fits and Memelink, 2000; Verdonk et al., 2005; Malitsky et al., 2008; Nieuwenhuizen et al., 2015; Zhang et al., 2015; Ying et al., 2020; Bomal et al., 2013; El-Azaz et al., 2020). Besides transcriptional reprogramming, primary metabolism enzymes can undergo biochemical changes that increase metabolic flux in the pathway it belongs to, allowing abundant and rapid production of downstream natural product(s). Consequently, some primary metabolism enzymes, especially at the interface of downstream natural product pathways, exhibit lineage-specific alterations in their feedback regulation (Bohlmann et al., 1996; Ning et al., 2015; Lopez-Nieves et al., 2018; Sugimoto et al., 2021; El-Azaz et al., 2022; Wang et al., 2022b). This coordinated regulation of primary and specialized metabolism in plants poses an intriguing question: if regulatory adjustment in primary metabolism evolved to ’support’ the new demand of emerging natural product pathways or, instead, if the downstream pathways emerged ’because of’ alteration in primary metabolism (Maeda, 2019; Yokoyama et al., 2024). Here, we used the abundance of genomic resources within grasses and other closely related Poales species to explore this question by investigating the evolutionary history of deregulated DHS1b and TyrAnc enzymes (**Figure 1a**), and their possible link to the emergence of the PTAL-mediated tyrosine-derived lignin pathway (**Figure 1b**).

The characterization of TyrAnc orthologs from Poales species outside of the core of the grass family (BOP and PACMAD clades, **Figure 1b**) indicates that deregulated TyrAnc was likely an ancestral trait that already existed in the common ancestor of Poales. This is supported by *in vitro* (**Table 1**, **Figure 3b**) and *in vivo* (**Figure 3c**) findings using AcTyrAnc from *Ananas comosus* within the family Bromeliaceae that is sister to the rest of Poales (McKain et al., 2016) and had its last shared ancestor with grasses between 125 to 190 million years ago (Givnish et al., 2018; Ma et al., 2021; Zhang et al., 2024). As bromeliad species are devoid of a PTAL ortholog and hence tyrosine-derived lignin biosynthesis (Takeda-Kimura et al., 2024), this implies that the original function of deregulated TyrAnc enzyme was not directly linked to the tyrosine-derived lignin pathway. This suggests that the emergence of the PTAL-mediated tyrosine-derived lignin pathway may have been a consequence of pre-existing high tyrosine levels, rather than the opposite. The TyrA phylogeny (**Figure 2**) also shows that plastidial TyrAnc is present in many monocot groups beyond Poales, though it remains to be examined if these TyrAnc enzymes are also deregulated. A recent study showed the high occurrence of tyrosine-derived specialized metabolites within some monocot lineages, such as in Amaryllidaceae and Liliaceae (Busta et al., 2024). Whether TyrAnc may be associated with the production of tyrosine-derived specialized metabolites in other monocot groups constitutes an alluring topic for future exploration.

Although TyrAnc enzymes from non-grass Poales are deregulated (**Table 1**), TyrAnc from grasses tend to have higher *K*i values (i.e., less sensitivity to tyrosine inhibition) (**Table 1**) and produce more tyrosine in *Nicotiana benthamiana* (**Figure 3c**). This tendency seems to be particularly apparent in TyrAnc enzymes from PACMAD grasses, such as SbTyrAnc and ZmTyrAnc (**Table 1**, **Figure 3c**). Synteny analysis of *TyrAnc* (**Supplemental Figure S1**) indicates this gene is within a conserved synteny block in grasses and other graminids, supporting their common origin and function. All things considered, deregulation of TyrAnc enzyme was likely present before, but possibly selected upon, the emergence of PTAL and the tyrosine-derived lignin pathway of graminids.

Phylogenetic analysis of the DHS family in Poales indicates that the deregulated DHS1b likely derived from the duplication of a single copy Poales *DHS1* gene (**Figure 4**). Although *in vitro* characterization of the DHS1a and DHS1b enzymes from *Pharus lappulaceous* revealed that PlDHS1b has lost sensitivity to inhibition by AAAs and arogenate (**Figure 5a**), *in planta* data show that transient expression of *PlDHS1b* does not trigger accumulation of high levels of AAAs or their pathway precursor molecules (**Figure 5b**). This finding suggests that the absence of sensitivity to inhibition by AAAs or arogenate *in vitro* is not necessarily a good predictor of DHS performance *in planta.* This may be due to the high diversity of effector molecules that can inhibit DHS enzymes through allosteric regulation, especially in plants (Webby et al., 2010; Jiao et al., 2012; Yokoyama et al., 2021; 2022a; 2022b).

*In planta* transient expression of DHSs from four different grass species show that only BdDHS1b and OsDHS1b, from *Brachypodium distachyon* and *Oryza sativa*, respectively, can cause a drastic metabolic effect (**Figures 5b**, **6**), which was not the case in the DHSs from the PACMAD grasses *Setaria viridis* and *Sorghum bicolor* (**Figure 6**). Thus, deregulation of DHS1b occurred within the grass family, more specifically in BOP grasses, which may confer these species certain adaptive advantage(s) by having a high flux shikimate pathway that can support high and rapid production of AAA-derived metabolites in varying environmental scenarios. Evolution of deregulated DHS1b within BOP species stays in clear contrast with deregulated TyrAnc that are widespread within Poales and might have allowed the grass ancestor to produce high levels of tyrosine-derived metabolites. Considering that PACMAD grasses lack the deregulated DHS but have the most deregulated TyrAnc enzymes (**Table 1**, **Figure 3c**), it is possible that PACMAD grasses preferentially produce lignin from tyrosine instead of phenylalanine.

The mutagenesis study of PlDHS1b, replacing selected residues by those of BdDHS1b, shows that high *in planta* DHS activity is caused by multiple mutations around the allosteric regulatory domain (**Figures 7d-h**), which corresponds to the effector binding site for tryptophan in MtDHA7P (Jiao et al., 2012). This supports a conserved function for this allosteric regulatory domain from bacteria to plant DHSs, although the exact effector molecule(s) regulating grass DHSs in *Nicotiana benthamiana* leaves remains unidentified. Some of these mutations, notably Thr211Met, are located within the *sota* domain (**Figure 7f**), a hot spot for mutations that deregulate Arabidopsis DHSs and cause drastic increases in AAAs levels in the plant (Yokoyama et al., 2022a). The Thr211Met substitution seems to be a major contributor to DHS de-regulation, whose impact becomes much stronger in combination with Ile262, Thr264, and Thr265 (**Figure 7h**), generating a DHS that is more active than the naturally de-regulated BdDHS1b. However, Thr211Met substitution is not present in OsDHS1b (**Supplemental Figure S10**) despite this protein still being highly active *in planta* (**Figure 6**). Therefore, our extensive mutagenesis analyses highlight that the alterations in multiple residues around the effector binding pocket synergistically modulate DHS feedback regulation, conveying that deregulated DHSs may evolve relatively easily in a convergent manner.

In summary, the comparative study of TyrA and DHS enzyme regulation across Poales and grasses revealed marked differences in the evolutionary timing of deregulated TyrAnc and DHS1b in relation to that of PTAL and the tyrosine-derived lignin pathway. Previously, altered feedback regulation of primary metabolism enzymes has been linked to the evolution of downstream specialized metabolism (Ning et al., 2015; Lopez-Nieves et al., 2018; Coley et al., 2019; Sugimoto et al., 2021; El-Azaz et al., 2022; Wang et al., 2022b). Our study suggests that increased precursor supply, due to the altered regulation of primary metabolism (e.g., deregulated tyrosine biosynthesis), may contribute to the emergence of novel specialized pathways (e.g., tyrosine-derived lignin biosynthesis), which can be subsequently fine-tuned by an additional alteration of primary metabolic regulation (e.g., DHS deregulation to enhance overall carbon flux). These findings highlight the dynamic and coordinated evolution of the primary and specialized metabolic interface, providing new insights and tools to rationally engineer plant metabolism to enhance the production of valuable chemicals.

## MATERIALS AND METHODS

### Plant material and growth conditions

The following grass cultivars were used a source for gene cloning: *Brachypodium distachyon* line 21-3, *Sorghum bicolor RTx430*, *Zea mays B73*, *Oryza sativa Nipponbare*, and *Setaria viridis*. *Nicotiana benthamiana* plants used in transient expression experiments were grown in 3x3 inches pots under ∼200 µE light intensity, 12h/12h photoperiod and 22°C, and watered regularly with a 12:4:8 (N:P:K) plant nutritive solution (Miracle-Gro) at a 1:1000 dilution in Hoagland’s solution diluted to 1:10 with water. *Joinvillea ascendens* seeds were obtained from David H. Lorence (National Tropical Botanical Garden; Kalaheo, Hawaii) and grown at the greenhouse facility of the Department of Botany, University of Wisconsin - Madison. For germination, *Joinvillea* seeds were treated with water at pH 2.0 for 24 hours at 30°C in the dark. The fruit flesh was removed prior to germination on soil. *Streptochaeta angustifolia* (specimen ID 2869/1) and *Pharus lappulaceous* plants (specimen ID 2452/1) were part of the Botany Greenhouse Collection at the Department of Botany, University of Wisconsin-Madison.

### **Cloning of** *TyrA* **and** *DHS* **gene CDSs**

*TyrA* and *DHS* coding sequences were cloned without the putative transit peptide (predicted using TargetP v2.0 server, DTU Health Tech) using the corresponding primers listed in **Supplemental Table S1**. All TyrA genes were cloned within the vector pET28a between the *Nde*I and *Bam*HI sites by In-Fusion cloning (Takara Bio, San Jose, California), or alternatively into the Golden Gate level 0 vector pAGM1287 (Weber et al., 2011). *TyrA1* genes from core grass species and *AcTyrA1* from *Ananas comosus* were directly cloned from genomic DNA, as these genes lack introns. Grass *TyrAnc* genes were all cloned from cDNA. *JaTyrA1, JaTyrAnc*, *SaTyrA1* and *SaTyrAnc* were cloned from cDNA. After various unsuccessful attempts of cloning pineapple’s *AcTyrAnc* from cDNA, we cloned the genomic version of this gene to confirm its sequence prior to synthesizing the CDS (Synbio Technologies, Monmouth Junction, NJ).

*DHS* genes were cloned from cDNA of the corresponding species into a modified version of the vector pET28a having *Bsa*I sites for Golden Gate sub-cloning, except *SbDHS1b, SbDHSnc* and *OsDHS1b*, which were synthesized into the vector pET100 D-TOPO (GeneArt, Thermo- Fisher). *Pharus lappulaceous PlDHS1a* and *PlDHS1b* gene sequence, without predicted plastid transit peptide, are available in GeneBank under the references PV055691 and PV055692, respectively.

Plant total RNA used for cloning was extracted from young leaf tissue using the CTAB/LiCl method based on Liao et al., 2004, with the modifications described in Canales et al., 2012. cDNA was synthesized with SuperScript IV VILO Master Mix (Thermo Scientific) following manufacturer’s instructions. Cloning PCRs were conducted using high fidelity DNA polymerase (PrimeSTAR Max DNA polymerase, Takara Bio, San Jose, California). PCR amplicons were purified from gel using QIAquick gel extraction kit (QIAGEN). All cloned genes were confirmed by Sanger sequencing.

### Site-directed mutagenesis and structural analysis of PlDHS1b

The PlDHS1b mutated residues grouped into the mutant blocks MUT1, MUT2 and MUT3, having a length of 700, 475 and 384 bp, respectively, are shown in **Supplemental Figure S9**. The three blocks were synthesized as gBlocks Gene Fragments (Integrated DNA Technologies, Coralville, Iowa, USA) overlapping 15 nucleotides between them at the sequences: 5’- ACCACAGCGAGCAGG (between gBlocks MUT1 and MUT2) and 5’-TGTCACCTGGGTCAC (between gBlocks MUT2 and MUT3). Block MUT1 DNA sequence had to be optimized between codons 16 and 41, both included (counting from the starting Met codon of the PlDHS1b sequence without predicted plastid transit peptide), to reduce %GC content, by using the online tool https://en.vectorbuilder.com/tool/codon-optimization.html (VectorBuilder, Chicago, Illinois). Mutant gBlocks were assembled altogether into *PlDHS1b^MUT1+2+3^* or, alternatively, with the fragments having the wild type *PlDHS1b* sequence, to produce *PlDHS1b^MUT1^*, *PlDHS1b^MUT2^* and *PlDHS1b^MUT3^*. Wild type *PlDHS1b* blocks shared same length as the synthesized mutant gBlocks and were made by amplifying these regions on *PlDHS1b* by PCR using specific primers listed in **Supplemental Table S1**. Blocks were assembled and cloned into a modified pET28a vector having flanking *Bsa*I sites for Golden Gate sub-cloning using In-Fusion cloning (Takara Bio, San Jose, California). *PlDHS1b^MUT4^* and *PlDHS1b^MUT2+3+4^* were generated by site-directed mutagenesis on pET28a-*PlDHS1b* or pET28a-*PlDHS1b^MUT2+3^*, respectively. *PlDHS1b^I262V_T264S_T265S^* and *PlDHS1b^T336A_E337D_R340K^* were generated by site-directed mutagenesis on pET28a-*PlDHS1b*. Sextuple mutant *PlDHS1b^M134T_T211M_I220V_I262V_T264S_T265S^* was generated by site-directed mutagenesis on pET28a-*PlDHS1b^I262V_T264S_T265S^.* All site-directed mutagenesis PCRs were done using high fidelity DNA polymerase (PrimeSTAR Max DNA polymerase, Takara Bio, San Jose, California). Primers used for site-directed mutagenesis are listed on **Supplemental Table S1**.

*In silico* homology modeling of PlDHS1b protein sequence was conducted using SWISS- MODEL (https://swissmodel.expasy.org/) (Waterhouse et al., 2018) under default parameters. The crystal structure of the tetrameric DHS enzyme from *Mycobacterium tuberculosis* (Protein Data Bank structure 3RZI, named MtDAH7P; Jiao et al., 2012) bound to its allosteric effector molecules, phenylalanine and tryptophan, was used as template. Representation of PlDHS1b homology model was done in CueMol2 Version 2.2.3.443 (http://www.cuemol.org/en/).

### Sequence identification and phylogenetic and synteny analyses

Plant species, genome version and source used for the TyrA and DHS protein sequences phylogenies (**Figures 2** and **4**) are listed in **Supplemental Table S2**. TyrA and DHS phylogenies were constructed by first obtaining the respective orthogroup using OrthoFinder version 2.5 (Emms and Kelly, 2019) with the default 1.5 MCL inflation parameter. After obtaining TyrA and DHS orthologs from OrthoFinder (orthogroups OG0001859 and OG0001685, respectively), sequences were filtered using Python scripts to remove duplicates, partial duplicates, and any gene with a length less than two times the standard deviation of the average length of all orthologs. Sequences were then aligned using MAFFT version 7 (https://mafft.cbrc.jp/alignment/server/; Katoh et al., 2019). ModelTest-NG was used to find the best evolutionary model for the data (Darriba et al., 2020). Finally RAxml-ng (Kozlov et al., 2019) was used to build the trees with the following parameters: For TyrA: --model JTT+I+G4+F --seed 210325 --tree pars{10},rand{10},OG0001859_tree.txt_mod2.txt --bs-metric fbp,tbe --bs-trees 200 --outgroup evm_27.model.AmTr_v1.0_scaffold00027.113,evm_27.model.AmTr_v1.0_scaffold00069.98; For DHS: --model JTT+I+G4+F --seed 210325 --tree pars{10},rand{10},OG0001685 _tree.txt_mod.txt --bs-metric fbp,tbe --bs-trees 200 --outgroup evm_27.model.AmTr_v1.0_scaffold00002.141,evm_27.model.AmTr_v1.0_scaffold00008.219 where –model is the model selected by ModelTest-NG, --seed is the random seed number, -tree has 10 parsimonious starting trees, 10 random starting trees, and one user input tree that was derived from the resolved gene tree from OrthoFinder, --bs-metric has both FBP and TBE bootstrap methods, --bs-trees means 200 bootstraps were used to calculate the bootstrap value, --outgroup indicates the Amborella genes used as outgroup. Python scripts and methods for running the various programs can be found in the following link: https://github.com/bmmoore43/Building-gene-trees.

TyrAnc synteny analysis was performed across selected Poales species using COGE (https://genomevolution.org/coge/) with the following options: DAGchainer: relative gene order, maximum distance between 2 matches: 20 genes, minimum number of aligned pairs: 5 genes, merged syntenic blocks with quota align, window size 100 genes, use all genes in the target genome, calculate synonymous substitution rates, and tandem duplication distance is 10. Synteny was calculated between the following species: *B. distachyon* (version 556, 3.0) and *J. ascendens* (version 1.1); *B. distachyon* (version 556, 3.0) and *P. latifolius* (version 1.0); *B. distachyon* (version 556, 3.0) and *S. bicolor* (version 454, 3.0.1); *P. latifolius* (version 1.0) and *S. angustifolia* (version 1.1); *S. angustifolia* (version 1.1) and *J. ascendens* (version 1.1); *J. ascendens* (version 1.1) and *P. latifolius* (version 1.0).

### Protein expression and purification

TyrA and DHS proteins were produced as N-terminus 6xHis tagged proteins in *Escherichia coli* strain KRX (Promega Corp.) using Terrific Broth (TB) medium, and purified using PureProteome Nickel Magnetic Beads (Millipore), as described by El-Azaz et al., 2023. Purified proteins were then buffer-exchanged into the corresponding TyrA or DHS storage buffer using a column pre-packed with Sephadex G-50 resin (GE Healthcare). For TyrA proteins, storage buffer consisted of 50 mM 4-(2-hydroxyethyl)-1-piperazineethanesulfonic acid (HEPES) buffer pH 7.5, 50 mM KCl, 10% glycerol, and 1 mM dithiothreitol (DTT). For DHS proteins, 50 mM HEPES buffer pH 7.5 supplemented with 300 mM NaCl, 0.2% Triton X-100 and 10% glycerol was used. In the case of DHS proteins, keeping NaCl concentration above 150 mM and including 10% glycerol or more was critical to avoid DHS protein precipitation. Buffer-exchanged proteins were frozen immediately in liquid nitrogen and stored at -80°C. Concentration of total protein was determined using Bio-Rad Protein Assay Dye Reagent Concentrate (Bio-Rad). Purity level of the recombinant enzymes was determined in ImageJ (v1.52a) upon staining of SDS-PAGE gel with Coomassie Brilliant Blue R250. Enzymatic assays were carried out within no longer than 2 weeks of protein storage at -80 °C, although we found many TyrAs and DHSs to be largely stable for longer periods (few months) in the storage buffers described.

### TyrA and DHS enzymatic assays

TyrA assays were conducted in a plate reader at 37 °C (Tecan Infinite M Plex, Tecan) using half-area plates (Greiner Bio-One) by tracking the conversion of NAD(P)^+^ into NAD(P)H as the increment of absorbance at 340 nm in 20 seconds intervals. TyrA reactions consisted of a final volume of 50 µL of 50 mM HEPES buffer pH 7.5, 50 mM KCl, 1 mM NADP^+^ (NAD^+^). The mass of TyrA enzyme per reaction was adjusted in between 10 to 200 ng, depending on the activity of the specific isoform, to ensure reaction’s linearity over time. Enzyme concentration was adjusted using TyrA desalting buffer supplemented with 0.5 mg/mL of Bovine Serum Albumin (BSA, protease-free powder purified by heat shock process; Fisher bioreagents). For *IC*50 assays, tyrosine was included into the reaction mixture pipetted from 10X-stocks adjusted to pH∼10 with NaOH, as tyrosine solubility is low at neutral pH. The reactions mixtures having the enzyme and all the reaction components, including tyrosine as inhibitor, but without substrate (arogenate) were incubated at 37 °C for 3 minutes before starting the reaction with the addition of substrate at 0.5 mM final concentration.

DHS activity was measured using a real-time method by tracking the consumption of phospho*enol*pyruvate at 232 nm (Schoner and Herrmann, 1976) at 37°C in a plate reader (Tecan Infinite M Plex, Tecan) in half-area UV-transparent 96-well plates (UV-Star^®^, Greiner Bio-One). DHS reaction consisted of a final volume of 50 µL of 25 mM HEPES buffer pH 7.5, 2 mM MgCl2, 3 mM DTT, the enzyme (variable mass, see details below), the effector molecule (AAAs or arogenate) and the substrates (phospho*enol*pyruvate and erythrose 4-phosphate). All DHS effectors tested were included in the initial reaction mixtures at a concentration of 0.5 mM. To ensure reactions linearity and good sensitivity, DHS mass per reaction was to 100-300 ng per reaction, depending on the specific activity of each isoform, using DHS storage buffer supplemented with 0.5 mg/mL of BSA. The reaction mixtures, having the enzyme and all the other components except phospho*enol*pyruvate or erythrose 4-phosphate, were incubated for 5 minutes at room temperature to allow the DTT-mediated activation of DHS. Then, phospho*enol*pyruvate was added to a final concentration of 1 mM, and an additional incubation step of 5 minutes at 37 °C was performed to stabilize absorbance at 323 nm. The enzymatic reaction was started with the addition of erythrose 4-phosphate at 1.5 mM final concentration.

Kinetic parameters of both TyrAs and DHSs were determined in MS-Excel using Solver add- in function. Arogenate was prepared by enzymatic conversion from prephenate (Prephenate Barium salt, Sigma-Aldrich) as previously described (Maeda et al., 2010).

### Plant expression constructs

For the transient expression in *Nicotiana benthamiana, TyrA* genes CDSs were amplified from pET28a constructs, without stop codon, and subcloned into the Golden Gate level 0 backbone pAGM1287 (Addgene plasmid #47996) by In-Fusion cloning (Takara Bio, San Jose, California). *TyrA* level 0 parts were then assembled into the Golden Gate binary vectors pAG4673 (for *PCaMV35S*-driven expression; Addgene plasmid #48014; Weber et al., 2011) or pICH47831 (for expression under control of *PSlRbcS3A*; Addgene plasmid #48010; Weber et al., 2011) as represented in **Supplemental Figure S3.** The plastid transit peptide from the enzyme 3-enol- pyruvyl-shikimate-3-phosphate synthase from *Petunia* × *hybrida* was used to target the TyrA enzyme into the plastid (Della-Cioppa et al., 1986). EU-NbHSP terminator was used in pAGM4673 constructs (Diamos and Mason, 2018). *YPet* was assembled to the C-terminus of *TyrA* genes for *in planta* protein stabilization. All other Golden Gate parts used are part of the MoClo Plant Parts Kit (Addgene Kit # 1000000047; Engler et al., 2014).

For *DHS* transient expression *in planta*, *DHS* CDSs, already cloned into Golden Gate compatible modified pET28a vector without plastid transit peptide or stop codon, were assembled into the Golden Gate binary vector pICH47822 (Addgene plasmid #48010; Weber et al., 2011) as represented in **Supplemental Figure S3.** *P*AtRbcS3B was used to drive *DHS* expression (MoClo Plant Parts Kit, Addgene Kit # 1000000047; Engler et al., 2014). The plastid signal transit peptide of RuBisCO (MoClo Plant Parts Kit, Addgene Kit # 1000000047; Engler et al., 2014) was used to direct DHS protein into the plastid.

Correct assembly of all Golden Gate parts into the plant expression plasmids was confirmed by colony PCR using specific primers targeting the promoter and terminator regions and digestion of the plasmid miniprep with restriction enzymes before transformation into *Agrobacterium tumefaciens* GV3101 by electroporation. *Agrobacterium tumefaciens* GV3101 transformant colonies were also tested by colony PCR using specific primers before proceeding to infiltration in *Nicotiana benthamiana*.

### **Transient expression in** *Nicotiana benthamiana*

*Agrobacterium tumefaciens* GV3101 clones transformed with the plant expression constructs were grown at 28°C for 24 to 36 hours in 3 mL of LB liquid media containing the corresponding antibiotics. The saturated cultures were spun down at 3,000 *g* for 5 minutes at room temperature and washed twice with 3 mL of induction media (IM; 10 mM MES [2-(N-morpholino)ethanesulfonic acid] buffer pH 5.6, 0.5% glucose, 2 mM NaH2PO4, 20 mM NH4Cl, 1 mM MgSO4, 2 mM KCl, 0.1 mM CaCl2, 0.01 mM FeSO4, and 0.2 mM acetosyringone). After the washing step, the cultures were incubated in IM for 2 to 3 hours at room temperature in the dark, followed by centrifugation at 3000 *g* for 5 minutes at room temperature, and resuspension into 3 mL of 10 mM MES buffer pH 5.6 supplemented with 0.2 mM acetosyringone. OD600nm was then adjusted to 0.25 (for pAGM4673-*TyrA* constructs) or 0.5 (for pICH47831-*TyrA* pICH47822-*DHS* constructs) prior to infiltration, using MES buffer with acetosyringone. pICH47831-TyrA and pICH47822-DHS clones were co-infiltrated with OD600nm∼0.5 of a p19 expressing construct to avoid gene silencing.

*Nicotiana benthamiana* plants of around 4-weeks-old were infiltrated close to the end of the light period into four different spots per plant, distributed into two leaves at two infiltrations per leaf according to a randomized pattern, with each individual spot corresponding to a different construct. Infiltrated *Nicotiana benthamiana* plants were kept under the same conditions as grown (see Plant material and growth conditions) until the end of the light period of the third day post-infiltration (∼72-hours post-infiltration). The infiltrated leaf tissue (leaf limbs without the main veins) was harvested and frozen immediately into liquid N2, ground to a fine powder and stored at -80 °C until analysis.

### Extraction of metabolites from plant tissue

Plant metabolites were extracted under alkaline pH following the procedure described in El-Azaz and Maeda, 2024. Around 30 to 40 mg of powder from deep frozen plant tissue were resuspended into 400 µL of cold chloroform:methanol mixed in 1:2 ratio and having 25 µM of L-norvaline, 0.5 µg/mL of isovitexin, and 10 µM of ^15^N-arogenate as internal standards. Extracts were kept for ∼1 hour on ice with regular vortexing, followed by centrifugation at 20,000 g for 5 minutes at room temperature. The supernantant was recovered and transferred to a fresh tube, then mixed with 300 µL of 1% 2-amino-2-methyl-1-propanol buffer at pH 10, followed with the addition of 150 µL chloroform. After 5 minutes of vigorous vortexing, the tubes were spun down at 20,000 g for 5 minutes at room temperature, and 250 µL of the aqueous supernantant were transferred twice into two independent tubes to be dried down overnight in a SpeedVac at ∼30°C. One of the dried pellets was stored as a backup at -20°C, the other resuspended into 100 µL of methanol 80% with 1 mM NaOH by sonication in a water bath at room temperature. After resuspension, the extracts were spun down at 20,000 g for 5 minutes at room temperature, and the supernantant transferred to vials for analysis.

### Quantification of plant metabolites

Tyrosine content in *Nicotiana benthamiana* upon *TyrA* transient expression was determined by HPLC (Infinity 1260, Agilent, Santa Clara, CA) equipped with a Water’s Atlantis T3 C18 column (3 μm, 2.1x150 mm) using in a gradient of water with 0.1% formic acid (mobile phase A) and acetonitrile with 0.1% formic acid (mobile phase B) at a flow rate of 0.3 mL/min and column temperature of 40 °C. The chromatographic gradient, 20 min duration in total, expressed as v/v % of mobile phase B in A, was: 0 to 5 min, 1% isocratic; 5 to 10 min, 1% to 76%; 10 to 12 min, 76% to 1%; 12 to 20 min, 1% isocratic. 10 μL of plant extract were used per injection. Tyrosine peak was detected at a retention time of ∼3.5 minutes using fluorescence detection mode (excitation wavelength 274 nm, emission wavelength 303 nm) and quantified with an authentic tyrosine standard (Alfa Aesar, catalog number AAA1114118).

Aromatic amino acids, and their pathway intermediates shikimate, prephenate and arogenate, were quantified using a Vanquish Horizon Binary UHPLC (Thermo Scientific) coupled to a Q-Exactive mass spectrometer (Thermo Scientific). One microliter of metabolite extract was injected onto a InfinityLab Poroshell 120 HILIC-Z column (150 × 2.1 mm, 2.7-μm particle size; Agilent) and resolved in a gradient of 5 mM ammonium acetate/0.2 % acetic acid buffer in water (phase A) and 5 mM ammonium acetate/0.2 % acetic acid buffer in 95% acetonitrile (phase B) at a flow rate of 0.45 mL/min and column temperature of 30 °C. Chromatographic gradients and MS settings (negative ionization mode) were as described in El-Azaz and Maeda, 2024. Spectral data were integrated manually using Xcalibur 3.0. Compound identity and abundance were determined based on calibration curves made using high purity authentic standards, and compound recovery was determined based on that of L-norvaline. All solvents used were LC-MS grade.

### Extraction of plant proteins and western blot

Total proteins from *Nicotiana benthamiana* samples were extracted from ∼10 mg of pulverized frozen tissue into 75 μL of 1X denaturing protein sample buffer (60 mM Tris [tris(hydroxymethyl)aminomethane] buffer pH 6.8, 2% sodium dodecyl sulfate, 10% glycerol, 3% β-mercaptoethanol, and 0.01% bromophenol blue) by vigorous vortexing for 30 seconds and boiled immediately at 95°C for 7 minutes. Tubes were centrifuged at 15,000 *g* for 5 minutes and 2.5 to 10 μL the supernatant, depending on the mass of plant tissue extracted and the accumulation level of the different TyrA or DHS, were applied to the SDS-PAGE gel. 3xFLAG tagged TyrA enzymes were detected using a monoclonal antibody conjugated to horseradish peroxidase (HRP) at a 1:1,000 dilution (OctA-Probe HRP conjugated antibody clone H-5, Sta. Cruz Biotechnology). TyrA-YPet fusion proteins were detected using a rabbit anti-GFP polyclonal antibody (A-6455, Invitrogen) at a 1:5,000 dilution and an anti-rabbit secondary antibody conjugated to HRP at a 1:10,000 dilution (SC-2357, Sta. Cruz Biotechnology). DHS-HA proteins were detected using a monoclonal antibody conjugated to HRP at a 1:1,000 dilution (HA-Probe HRP conjugated mouse monoclonal antibody clone F-7, Sta. Cruz Biotechnology, cat. no. SC- 7392). All antibodies were prepared at the indicated dilution in Tris-buffered saline 1X (20 mM Tris-HCl, 150 mM NaCl, pH 7.4) with 0.5% BSA (protease-free powder purified by heat shock process; Fisher bioreagents). The level of transgenic enzymes was measured by ImageJ (version 1.52a) comparing the chemiluminescent signal in the plant samples with a standard curve made of pure recombinant SbTyrA1-3xFLAG, SbTyrA1-YPet or BbDHS1b-3xHA protein produced in *E. coli*.

### **YPet imaging in** *Nicotiana benthamiana*

Infiltrated leaf areas expressing *TyrA*-*YPet* constructs were imaged with a Zeiss LSM710 confocal imaging system (Newcomb Imaging Center, Department of Botany, UW-Madison). Plant limbs were directly imaged at 72 hours post-infiltration with a 514 nm argon laser. Fluorescence was detected between 516 - 550 nm (for YPet) and 650 - 750 nm (for chlorophyll) with a Plan Apochromat 20x/0.8 objective. At least three representative regions from two to three independent infiltrated leaves were imaged. Images were processed using Zen software (Zeiss).

### Statistics and Reproducibility

All experiments shown in the manuscript were conducted at least twice to confirm the reproducibility of the findings. Statistical comparisons between samples (Student’s *t*-test) were done in Microsoft Office Excel. No data were excluded from any of the analyses shown. No statistical method was used to predetermine sample size. All the experiments conducted with plants were randomized. The Investigators were not blinded to allocation during experiments and outcome assessment.

## SUPPLEMENTAL DATA

The following materials are available in the online version of this article.

Supplemental Table S1. List of primers used in this study.

Supplemental Table S2. Sequence and species information.

Supplemental Figure S1. Synteny analysis of Poales TyrAnc.

Supplemental Figure S2. Michaelis-Menten plots corresponding to the kinetical parameters *K*m and *V*max for TyrA enzymes.

Supplemental Figure S3. Golden Gate constructs for transient expression in *Nicotiana benthamiana*.

Supplemental Figure S4. Transient overexpression of Poales *TyrAnc* genes in *Nicotiana benthamiana* with C-terminal 3xFLAG tag, under control of *CaMV 35S* promoter.

Supplemental Figure S5. Laser scanning confocal microscopy of *Nicotiana benthamiana* leaves expressing TyrA-YPet fusion proteins.

Supplemental Figure S6. Immunoblot of TyrAnc-YPet fusion proteins in *Nicotiana benthamiana*.

Supplemental Figure S7. Immunoblot of grass and Poales DHS-HA fusion proteins in *Nicotiana benthamiana*.

Supplemental Figure S8. Immunoblot of BdDHS1b, SbDHS1b, SvDHS1b and OsDHS1b HA fusion proteins in *Nicotiana benthamiana*.

Supplemental Figure S9. Multiple sequence alignment of deregulated BdDHS1b and OsDHS1b, compared to Arabidopsis DHSs.

Supplemental Figure S10. Multiple sequence alignment of DHS proteins from graminids.

Supplemental Figure S11. Transient overexpression of PlDHS1b with additional MUT4 mutations.

Supplemental Figure S12. Immunoblot of *Pharus lappulaceous* DHS1b wild type and mutated proteins C-terminal 3xHA fusion proteins in *Nicotiana benthamiana*.

## ACKNOWLEDGEMENTS

We are thankful to our collaborators James Leebens-Mack (Department of Plant Biology, University of Georgia) and Michael McKain (Department of Biological Sciences, University of Alabama) for obtaining the *Joinvillea ascendens* genome sequence, Matthew Moscou (USDA-ARS, Cereal Disease Laboratory at St. Paul) for sharing the *Ecdeiocolea monostachya* genome, Ryo Yokoyama (UW-Madison) for the original *SbDHS1b*, *SbDHSnc*, *OsDHS1b* and *OsDHSnc* constructs, and Yuri Takeda-Kimura (Yamagata University, Yamagata, Japan) for critically reading the manuscript draft. The seeds of *Joinvillea ascendens* were a courtesy of David H. Lorence (National Tropical Botanical Garden; Kalaheo, Hawaii). YPet fluorescent protein was kindly provided by Dr. Makoto Yanagisawa (UW-Madison). Golden Gate vectors and plant modular cloning parts are courtesy of Sylvestre Marillonnet (Leibniz-Institut für Pflanzenbiochemie; Halle, Sachsen-Anhalt, Germany) and Nicola Patron (The Earlham Institute; Norwich, United Kingdom). This work was supported by the U.S. National Science Foundation (NSF) PGRP-IOS-1836824 and MCB-2404174 to H.A.M.

## Author’s Contribution

H.A.M. and J.E.A. planned and designed the research. J.E.A. conducted the experiments. B.M. performed phylogenetic analysis and ortholog identification of TyrA and DHS families. J.E.A. and H.A.M. wrote the manuscript with input from B.M.

## Conflicts of interest statement

HAM and JEA have filed for a patent related to the naturally deregulated enzyme BdDHS1b (described by El-Azaz et al., 2023) and the newly identified OsDHS1b reported in this study (Publication number: 20250115926, filed on October 4, 2024). The authors declare that they have no other competing interests.

**Table 1.** Kinetic parameters of grasses and non-grass poales TyrA enzymes. Kinetic parameters *K*m and *V*max were obtained from the Michaelis–Menten curves obtained by TyrA activity measured at various concentrations of arogenate (Supplemental Figure S1). *k*cat was deduced from *V*max and the molecular weight of each recombinant enzyme, including the mass of poli-histidine tag. Apparent sensitivity to inhibition by tyrosine (*IC*50) was determined at variable concentrations of tyrosine at a fixed concentration of 500 µM of arogenate and 1 mM of NADP^+^. The inhibition constant for tyrosine (*K*i) was calculated from *K*m and *IC*50 values under a competitive inhibition model. All data are means ± *SD* of *n* ≥ 2 independent experiments conducted on different days with different batches of purified enzyme. Enzymes marked with *** were previously characterized by El-Azaz et al., 2023 under these same conditions and have been included for comparison purposes.

| | | $K_m$<br>(μM Agn) | $V_{max}$<br>(pKat/μg) | $k_{cat}$<br>(s<sup>-1</sup>) | $k_{cat}/K_m$<br>(s<sup>-1</sup>·mM<sup>-1</sup>) | $IC_{50}$<br>(μM Tyr) | $K_i$<br>(μM Tyr) |
| --- | --- | --- | --- | --- | --- | --- | --- |
| BdTyrA1<br>orthologs | AcTyrA1 | 750 ± 80 | 919 ± 186 | 33 ± 7 | 43 ± 9 | 20 ± 10 | 12 ± 6 |
| | JaTyrA1 | 730 ± 60 | 516 ± 27 | 18 ± 1 | 25 ± 1.3 | 40 ± 7 | 24 ± 4 |
| | SaTyrA1 | 720 ± 90 | 86 ± 1 | 3.2 ± 0.03 | 4.4 ± 0.05 | 57 ± 38 | 34 ± 23 |
| | OsTyrA1 | 320 ± 40 | 84 ± 5 | 3.0 ± 0.2 | 9.4 ± 0.6 | 53 ± 5 | 21 ± 2 |
| | ZmTyrA1a | 530 ± 50 | 143 ± 3 | 5.2 ± 0.1 | 9.7 ± 0.2 | 46 ± 5 | 24 ± 2 |
| | BdTyrA1*** | 1470 ± 240 | 134 ± 16 | 5.2 ± 0.6 | 3.5 ± 0.4 | 66 ± 2 | 49 ± 2 |
| | SbTyrA1*** | 550 ± 130 | 87 ± 17 | 3.2 ± 0.6 | 5.8 ± 1.1 | 71 ± 24 | 37 ± 13 |
| BdTyrAnc<br>orthologs | AcTyrAnc | 1050 ± 170 | 162 ± 8.1 | 6.4 ± 0.3 | 6.7 ± 0.3 | 110 ± 40 | 76 ± 30 |
| | JaTyrAnc | 1900 ± 240 | 2458 ± 218 | 98 ± 9 | 51 ± 5 | 130 ± 20 | 106 ± 19 |
| | SaTyrAnc | 350 ± 20 | 2408 ± 28 | 88 ± 1 | 251 ± 3 | 85 ± 20 | 35 ± 8 |
| | OsTyrAnc | 190 ± 50 | 3310 ± 132 | 119 ± 5 | 616 ± 25 | 280 ± 70 | 78 ± 19 |
| | ZmTyrAnc | 350 ± 70 | 7423 ± 423 | 293 ± 17 | 832 ± 47 | 850 ± 30 | 352 ± 13 |
| | BdTyrAnc*** | 140 ± 40 | 2036 ± 107 | 74 ± 4 | 515 ± 27 | 240 ± 40 | 54 ± 10 |
| | SbTyrAnc*** | 220 ± 60 | 1880 ± 475 | 73 ± 18 | 337 ± 85 | 470 ± 150 | 144 ± 47 |

