## Supplemental Figures File for "Dynamic Retuning of Precursor Supply Before and After the Evolution of the Dual Lignin Pathway in Poales"

2025

**List of content**

**Supplemental Figure S1**. Synteny analysis of Poales *TyrA*$_{nc}$.

**Supplemental Figure S2**. Michaelis-Menten plots corresponding to the kinetical parameters $K_m$ and $V_{max}$ for TyrA enzymes.

**Supplemental Figure S3**. Golden Gate constructs for transient expression in *Nicotiana benthamiana*.

**Supplemental Figure S4.** Transient overexpression of Poales *TyrA*$_{nc}$ genes in *Nicotiana benthamiana* with C-terminal 3xFLAG tag, under control of *CaMV 35S* promoter.

**Supplemental Figure S5**. Laser scanning confocal microscopy of *Nicotiana benthamiana* leaves expressing TyrA-YPet fusion proteins.

**Supplemental Figure S6**. Immunoblot of TyrAnc-YPet fusion proteins in *Nicotiana benthamiana.*

**Supplemental Figure S7.** Immunoblot of grass and Poales DHS-HA fusion proteins in *Nicotiana benthamiana*.

**Supplemental Figure S8.** Immunoblot of BdDHS1b, SbDHS1b, SvDHS1b and OsDHS1b HA fusion proteins in *Nicotiana benthamiana*.

**Supplemental Figure S9**. Multiple sequence alignment of deregulated BdDHS1b and OsDHS1b, compared to Arabidopsis DHSs.

**Supplemental Figure S10**. Multiple sequence alignment of DHS proteins from graminids.

**Supplemental Figure S11**. Transient overexpression of PlDHS1b with additional MUT4 mutations.

**Supplemental Figure S12**. Immunoblot of *Pharus lappulaceous* DHS1b wild type and mutated proteins C-terminal 3xHA fusion proteins in *Nicotiana benthamiana*.

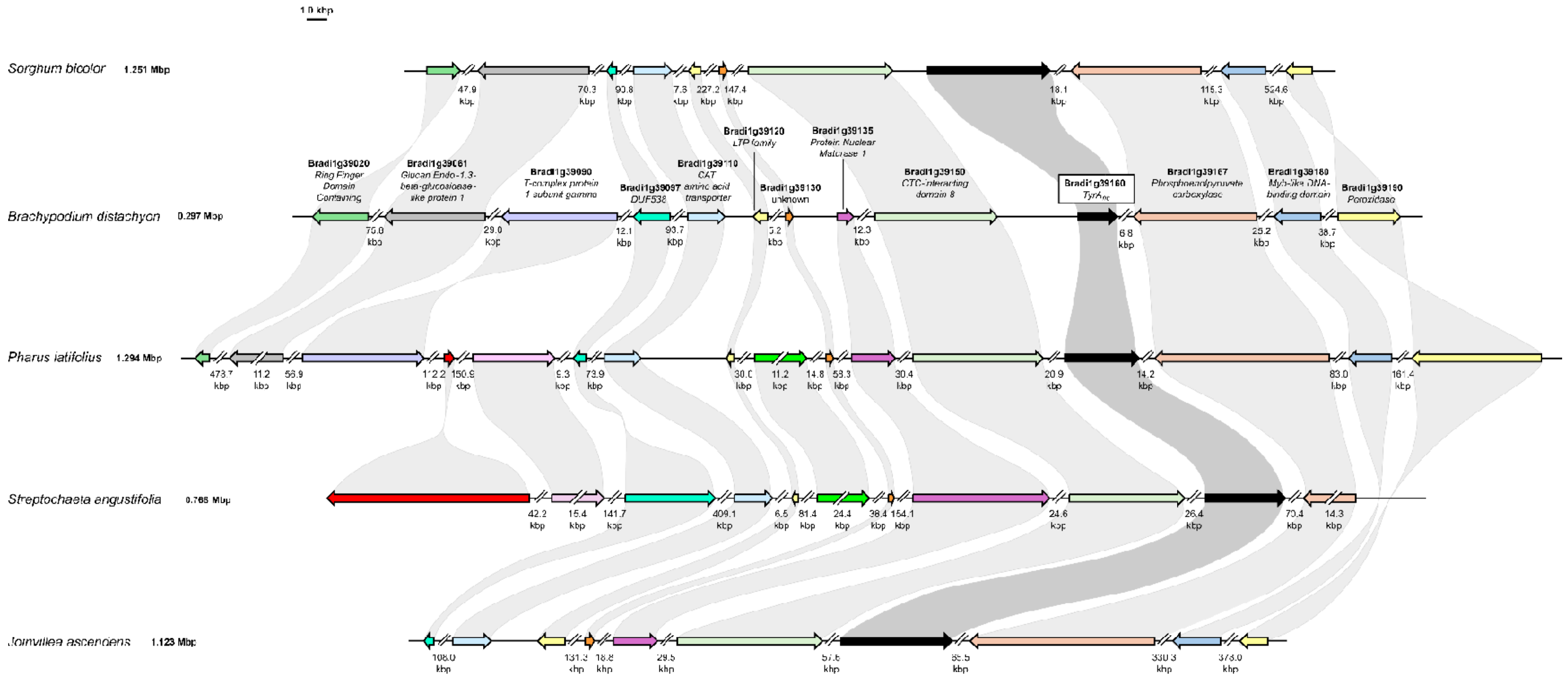
1.0 kbp
Sorghum bicolor 1.251 Mbp
Brachypodium distachyon 0.297 Mbp
Pharus latifolius 1.294 Mbp
Streptochaeta angustifolia 0.765 Mbp
Joinvillea ascendens 1.123 Mbp
Bradi1g39020 Ring Finger Domain Containing
Bradi1g39061 Glucan Endo-1,3-beta-glucosidase-like protein 1
Bradi1g39090 T-complex protein 1 subunit gamma
Bradi1g39097 DUF538
Bradi1g39110 CAT amino acid transporter
Bradi1g39120 LTP family
Bradi1g39130 unknown
Bradi1g39135 Protein Nuclear Maturase 1
Bradi1g39150 CTC-interacting domain 8
Bradi1g39160 TyrA
Bradi1g39167 Phosphoenolpyruvate carboxylase
Bradi1g39180 Myb-like DNA-binding domain
Bradi1g39190 Peroxidase

**Supplemental Figure S1. Synteny analysis of Poales *TyrA$_{nc}$* (previous page).** The análisis, performed using CoGe database (see Methods), shows TyrAnc genes are in synteny across graminid species. Orthologous genes are highlighted with light gray shades. Double slashes denote genomic regions that are not shown in scale (size indicated below double slashes in kbp).

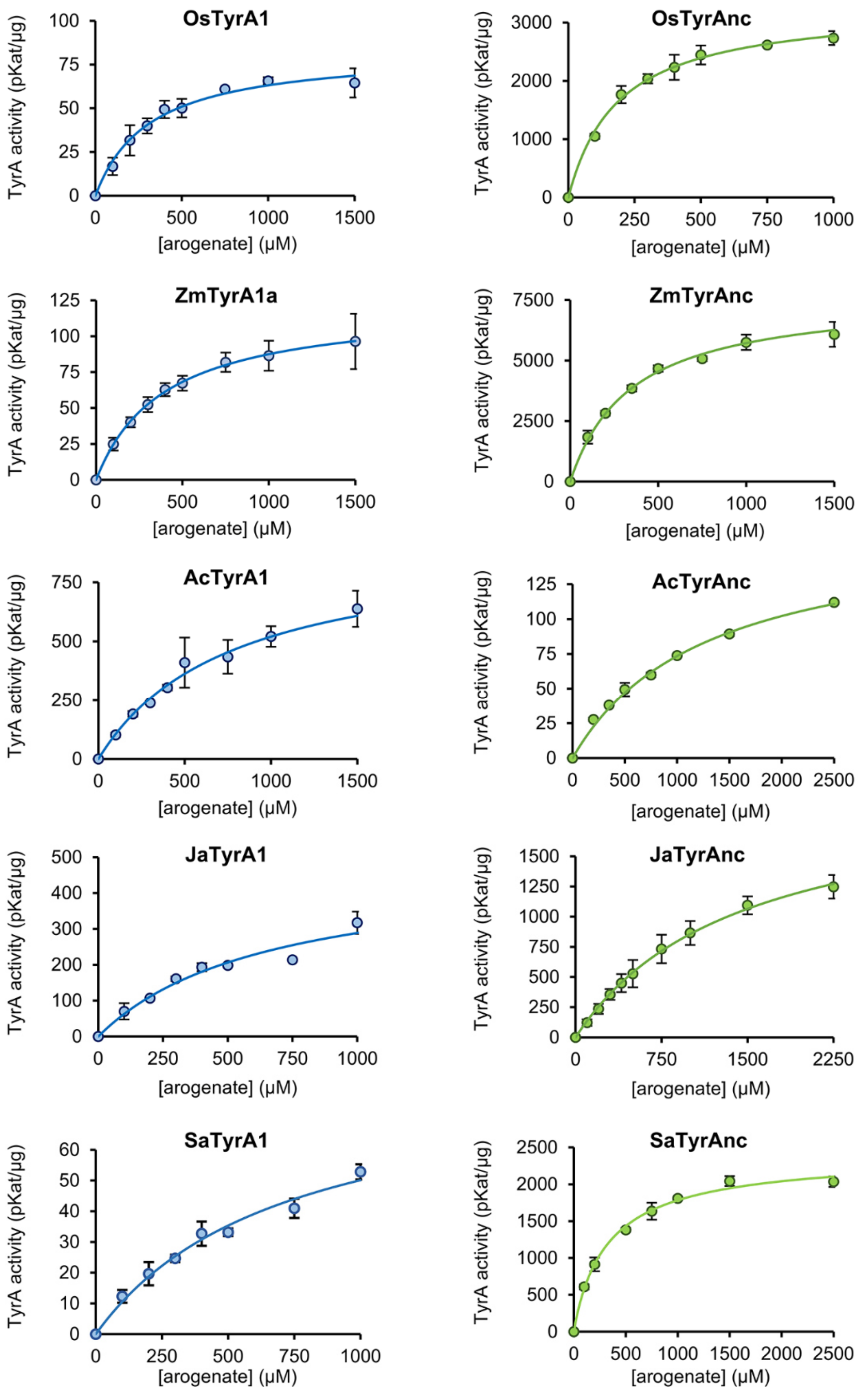


**Supplemental Figure S2**. **Michaelis-Menten plots corresponding to the kinetical parameters $K_m$ and $V_{max}$ for TyrA enzymes, as shown in main Table 1.** Individual points represent the average of at least two replicates from independent experiments, using different preparations of both purified recombinant enzyme and purified arogenate. Error bars = *SD*.

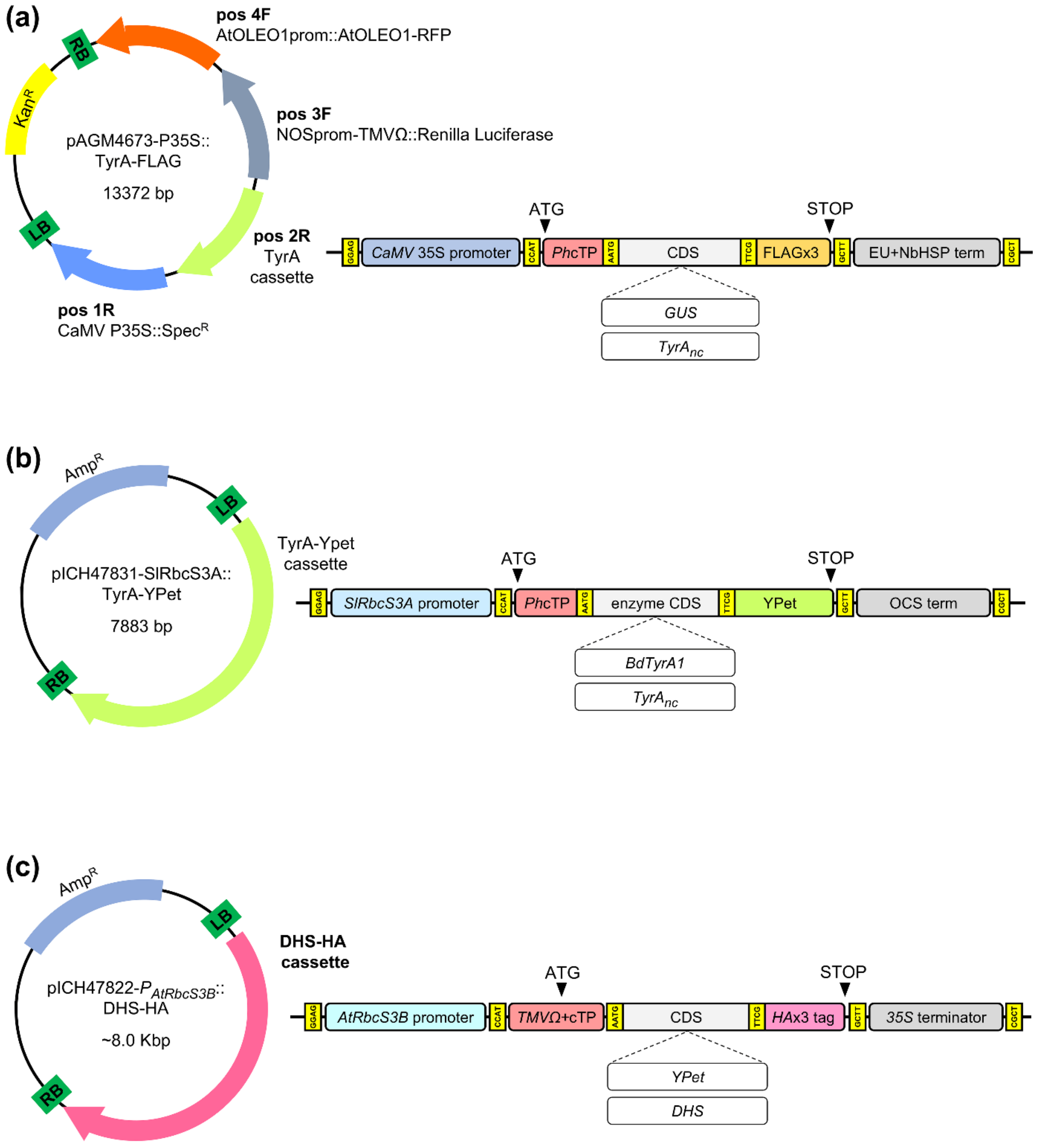


**Supplemental Figure S3. Golden Gate constructs for transient expression in *Nicotiana benthamiana.*** **(a)** *TyrA* expression under control of *CaMV* 35S promoter or **(b)** tomato's *SlRbcS3A* promoter. **(c)** *DHS* expression construct. Yellow boxes indicate Golden Gate overhangs used for the modules assembly. *Phc*TP, putative plastid transit peptide of *Petunia* x *hybrida* 5-enol-pyruvyl-shikimate-3-phosphate synthase (Della-Cioppa et al., 1986). *TMVΩ*+cTP, Tomato Mosaic Virus Ω enhancer fused to RuBisCO consensus plastid transit peptide (Engler et al., 2014). EU+NbHSP terminator was based on Diamos and Mason (2018). Plant Biotechnology Journal, 16:1971-1982. OCS term, *Agrobacterium tumefaciens* octopine synthase terminator.

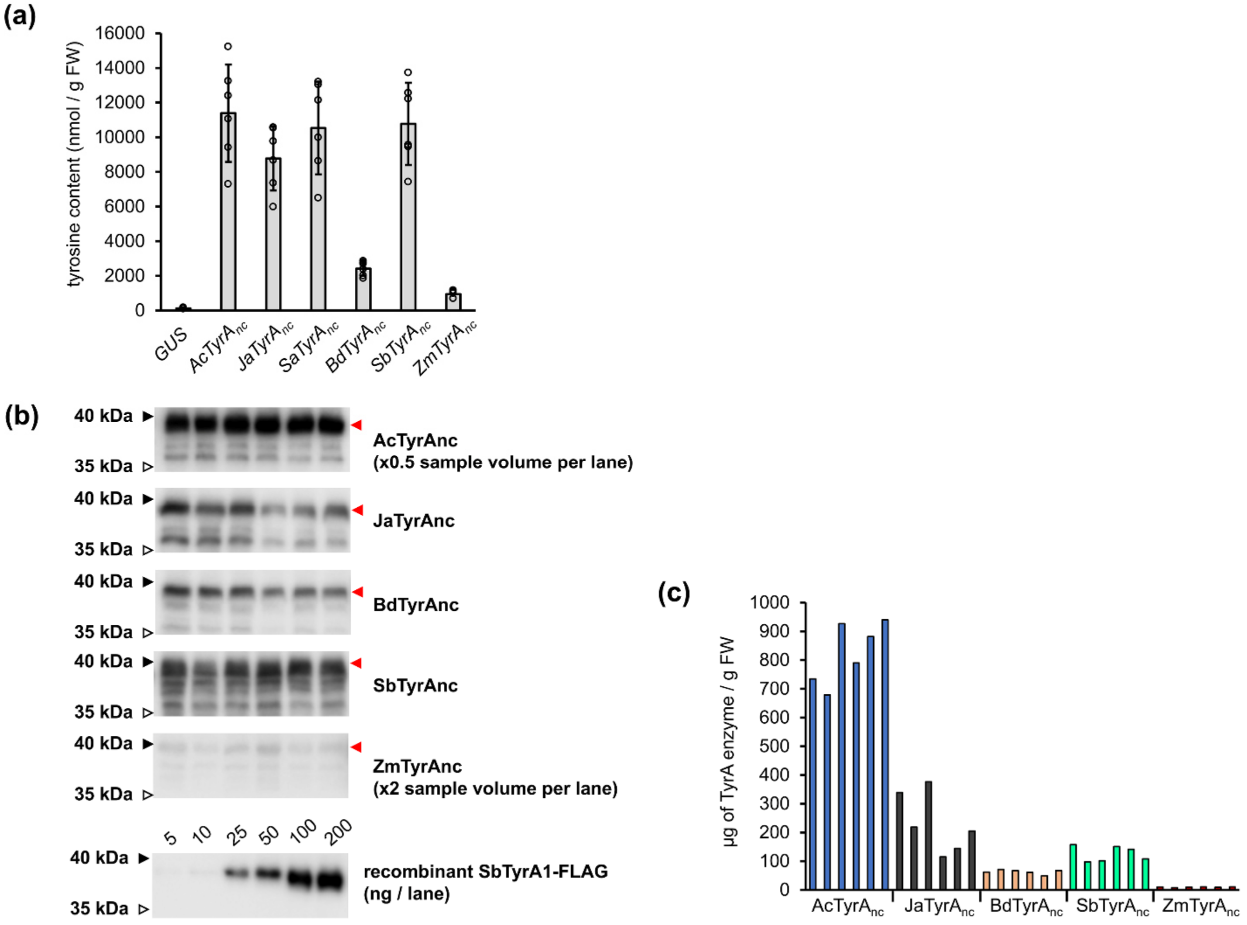


**Supplemental Figure S4. Transient overexpression of Poales *TyrA$_{nc}$* genes in *Nicotiana benthamiana* with C-terminal 3xFLAG tag, under control of *CaMV 35S* promoter. (a)** Tyrosine content in plant samples, determined by HPLC-FLD at 72 hours post-infiltration. Bar graphs represent the average of *n* = 6 biological replicates coming from independent plants. Error bars = *SD.* **(b)** Anti-FLAG tag immunoblot of total proteins extracted from plant samples and calibration curve using recombinant SbTyrA1 enzyme tagged with FLAG. The volume of sample per lane was reduced by half for AcTyrA$_{nc}$ due to the strong chemiluminescent blot signal observed in preliminary immunoblots. Sample volume per lane was duplicated for ZmTyrA$_{nc}$ samples due to the very low abundance of the protein. Red arrows indicate the expected size of the mature protein without transit peptide. All images correspond to the same western blot experiment. **(c)** TyrA protein abundance per individual plant sample, as estimated by image-based analysis of anti-FLAG tag immunoblot. Each bar graph represents a single biological sample coming from an independent plant.

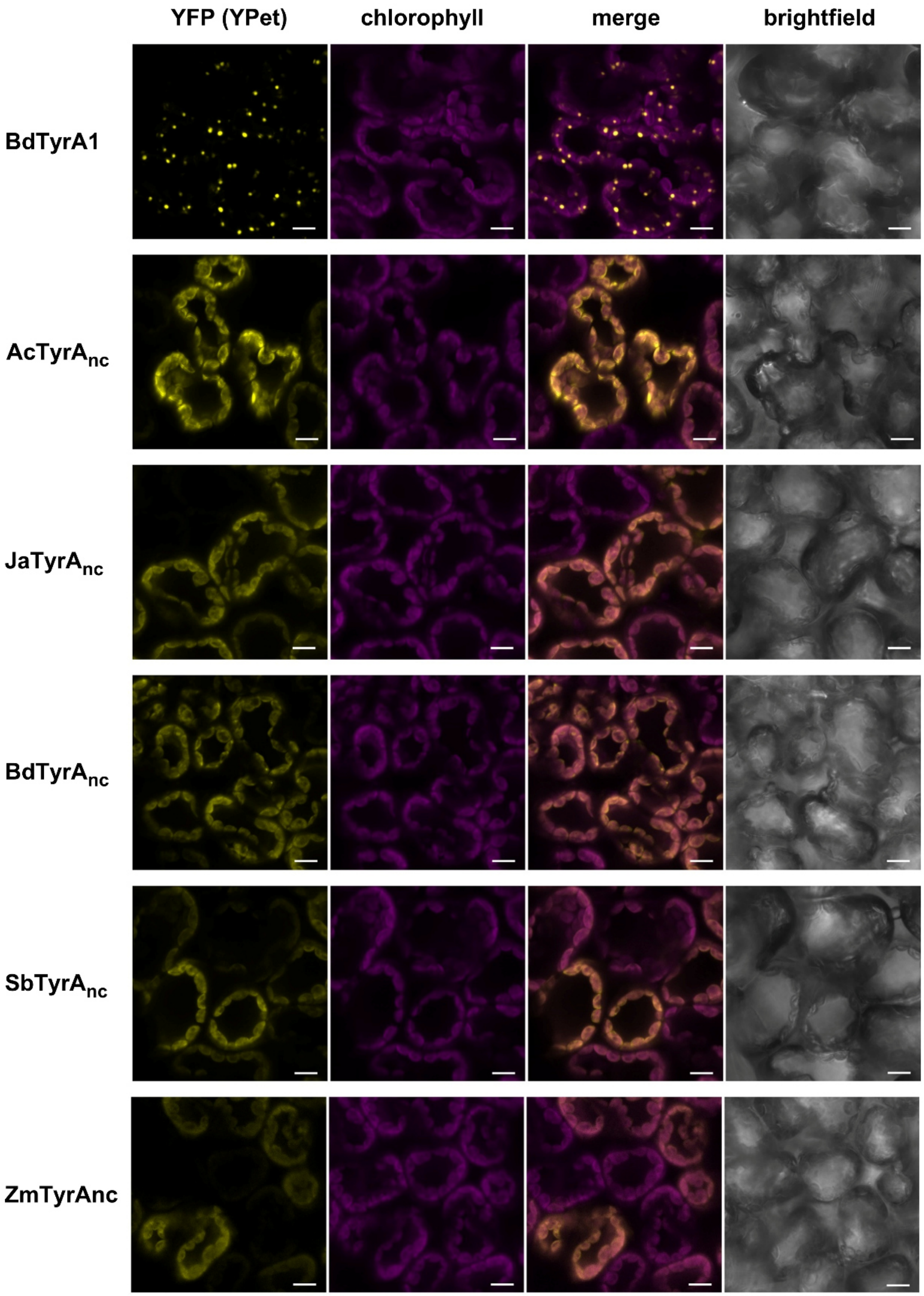


**Supplemental Figure S5**. **Laser scanning confocal microscopy of *Nicotiana benthamiana* leaves expressing TyrA-YPet fusion proteins.** Plants were imaged at 3 days post-infiltration. Z-projections shown are representative results across three independent infiltrated plants. Scale bar = 10 µm.

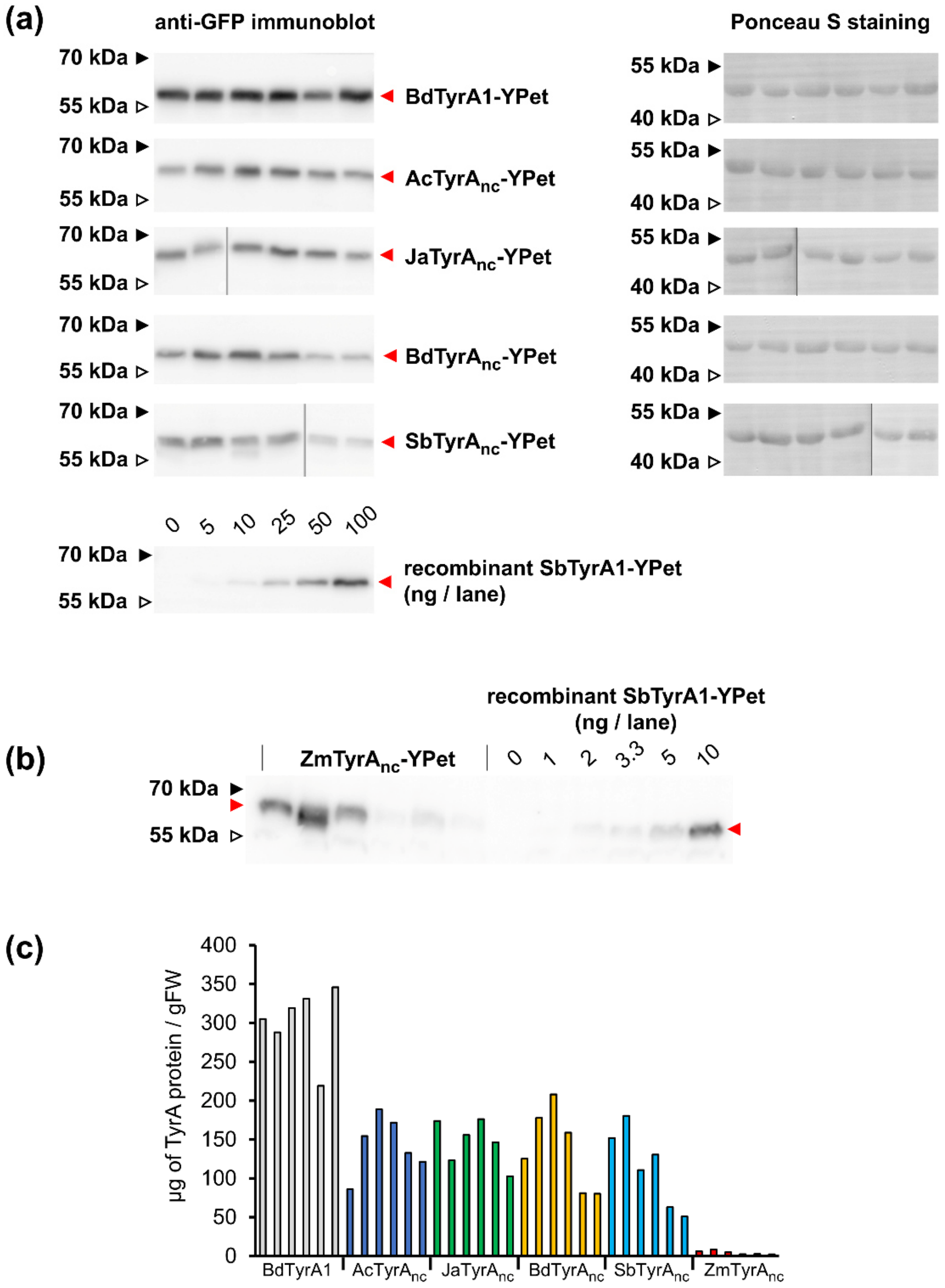


**Supplemental Figure S6**. **Immunoblot of TyrAnc-YPet fusion proteins in *Nicotiana benthamiana*. (a)** Anti-GFP immunoblot and on-membrane Ponceau S staining of total proteins extracted from plant samples and calibration curve using recombinant SbTyrA1-YPet enzyme. Red arrows indicate the expected size of the mature protein without transit peptide. All images correspond to the same western blot experiment. Vertical lines in gray color in $JaTyrA_{nc}$ and $SbTyrA_{nc}$ samples separate non-consecutive lanes. Bar graphs represent the average of *n* = 6 biological replicates coming from independent plants. Error bars = *SD.* **(b)** Anti-GFP immunoblot of total proteins in $ZmTyrA_{nc}$ samples and associated standard curve using SbTyrA1-YPet. This immunoblot is independent from the results shown in A, as the abundance of $ZmTyrA_{nc}$ was found to be much lower in preliminary immunoblots. **(c)** TyrA enzyme abundance per individual plant sample, as estimated by image-based analysis of anti-GFP immunoblots. Each bar graph represents a single biological sample coming from an independent plant.

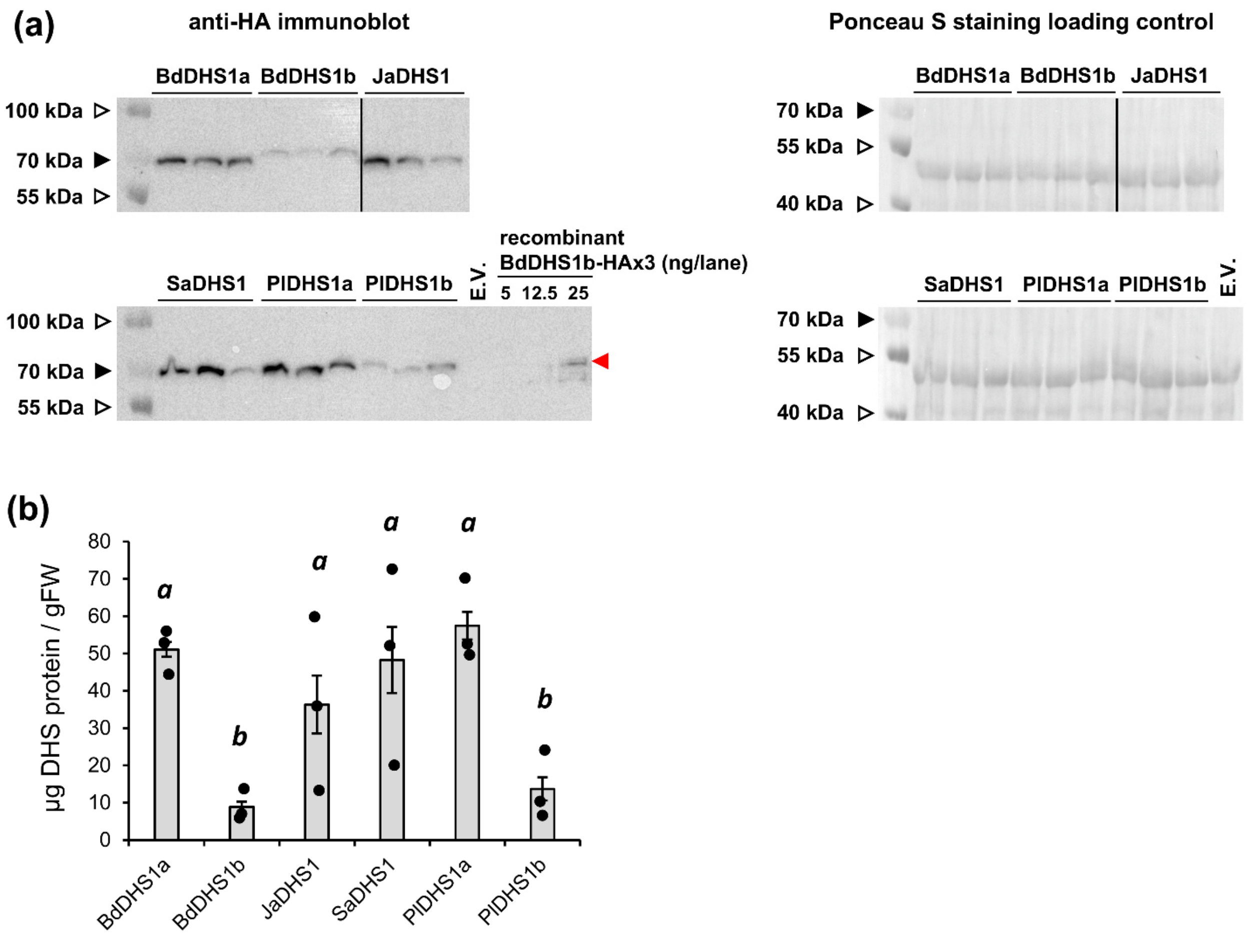


**Supplemental Figure S7**. **Immunoblot of grass and Poales DHS-HA fusion proteins in *Nicotiana benthamiana*. (a)** Anti-HA immunoblot and on-membrane Ponceau S staining of total proteins extracted from plant samples and calibration curve using recombinant BdDHS1b-HAx3 protein (mass in ng indicated above each lane; only the upper BdDHS1b-HAx3 band was used for quantification). Each lane corresponds to an independent biological sample. Red arrows indicate the expected size of the mature protein without transit peptide. All images correspond to two SDS-PAGE gels that were transferred to a single membrane and analyzed in the same western blot experiment. **(b)** Quantification of DHS abundance according to the results shown in (a) and normalized by mass of plant tissue extracted (*n* = 5; error bars = *SE*).

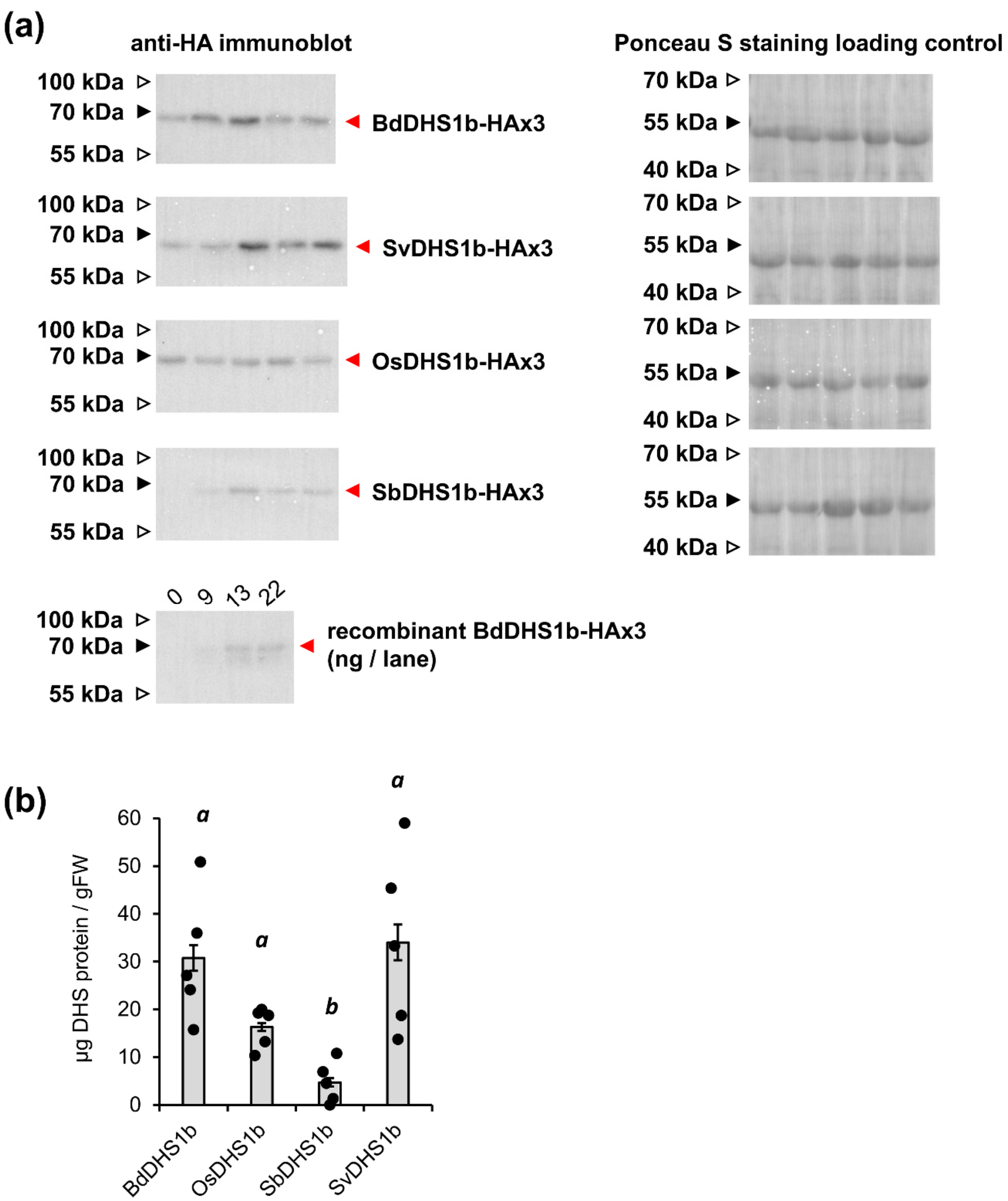


**Supplemental Figure S8. Immunoblot of BdDHS1b, SbDHS1b, SvDHS1b and OsDHS1b HA fusion proteins in *Nicotiana benthamiana*. (a)** Anti-HA immunoblot and on-membrane Ponceau S staining of total proteins extracted from plant samples and calibration curve using recombinant BdDHS1b-3xHA protein (mass in ng indicated above each lane; only the upper BdDHS1b-3xHA band was used for quantification). Each lane corresponds to an independent biological sample. Red arrows indicate the expected size of the mature protein without transit peptide. All images correspond to the same western blot experiment. **(b)** Quantification of DHS abundance according to the results shown in (a) and normalized by mass of plant tissue extracted ($n$ = 5; error bars = $SE$).

```
          1                                                                                                      100
OsDHS1b   AVHAAEPSKN PPVGVPSAAK TSSPSVAAPE KAPVAAAPAP VAPAPAATKQ VAPARWAVDS WRTKKALQLP EYPNAAELEA VLKTIEAFPP IVFAGEARHL
BdDHS1b   AVHAAEPSKS AAAA-PAAAK TSSPSVAPEK SAIPDPKPEA PAV------- --PAKWTVDS WRAKKALQLP EYPNAAELES ALKTIEAFPP IVFAGEARHL
PlDHS1b   AVHAAEPSKN HSSI-PAAAK TSSPTVAP-- ---------- -APAPEKAAD APPARWSVDS WRAKKALQLP EYPNAAELEA VLKNIEAFPP IVFAGEARHL
BdDHS1a   AVHAADPARG TGS-VPAAAK TSSPTVAPEA AASPA----- ---------- ---ASWKVDS WKAKKALQLP EYPSKEELDT VLQTIEAFPP IVFAGEARHL
 AtDHS3                       A VQTDPKTPAA SSASAATTTP ATLTKPVGVN VGKGKWAPES WRTKKALQQP DYPDLAALEA VLETIEAFPP IVFAGEARLL
 AtDHS1   AVHAAEPARN AVSVKESVAS SSS------- ---------- ---------- -GALKWTPES WKLKKALQLP DYPNANELES VLKTIEAFPP IVFAGEARNL
 AtDHS2   AVHSTDPKKS TQSASASV-- ---------- ---------- ---------- ----KWSLES WKSKKALQLP DYPDQKDVDS VLQTLSSFPP IVFAGEARKL

          101                                                                                                    200
OsDHS1b   EERLADAAMG RAFLLQGGDC AESFKEFNGN NIRDTFRVLL QMSAVLTFGG QMPVIKVGRM AGQFAKPRSE AFEERDGVKL PSYRGDNING DAFNEKSRIP
BdDHS1b   EERLADAAMG RAFLLQGGDC AESFKEFNGN NIRDTFRVLL QMSAVLTFGG QMPVIKVGRM AGQFAKPRSD SFEVRDGVKL PSYRGDNING DAFNEKSRIP
PlDHS1b   EERLADAAMG RAFLLQGGDC AESFKEFNAN NIRDTFRVLL QMSAVLMFGG QMPVVKVGRM AGQFAKPRSD HFEVRDGVKL PSYRGDNVNG DAFDEKSRVP
BdDHS1a   EERLAEAAMG RAFVLQGGDC AESFKEFNAN NIRDTFRVLL QMGAVLMFGG QVPVVKVGRM AGQFAKPRSD NLEERDGVKL PSYRGDNVNG DAFDVKSRTP
 AtDHS3   EERLGQAAMG EAFLLQGGDC AESFKEFNAN NIRDTFRILL QMGAVLMFGG QVPVVKVGRM AGQFAKPRSD SFEEKDGVKL PSYRGDNING DAFDSKSRIP
 AtDHS1   EERLADAAVG KAFLLQGGDC AESFKEFNAT NIRDTFRVLL QMSIVLTFGG QVPVIKVGRM AGQFAKPRSD AFEEKDGVKL PSYKGDNING DTFDEKSRIP
 AtDHS2   EDKLGQAAMG QAFMLQGGDC AESFKEFNAN NIRDTFRVLL QMGVVLMFGG QLPVIKVGRM AGQFAKPRSD PFEEKDGVKL PSYRGDNING DAFDEKSRIP

          201                   sota helix-loop-helix                                                            300
OsDHS1b   DPQRMVRAYA QSAATLNLLR AFATGGYAAM QRVTQWNLDF TQHSEQGDRY RELAHRVDEA LGFMSAAGLT VDHPLMTSTD FWTSHECLLL PYEQSLTRQD
BdDHS1b   DPQRMIRAYT QSAATLNLLR AFAMGGYAAM QRVTQWNLDF TENSEQGDRY RELAHRVDEA LGFMSAAGLT LDHPVMSSTE FWTSHECLLL PYEQALTRQD
PlDHS1b   DPQRMIRAYA QSAATLNLLR AFATGGYAAM QRITQWNLDF TDHSEQGDRY RELAHRVDEA LGFMAAAGLT VDHPIMTTTE FSTSHECLLL PYEQALTRED
BdDHS1a   DPERMIRAYA QSVATLNLLR AFATGGYAAM QRVIQWNLDF MDHNEQGDRY RELAHRVDEA LGFMTAAGLG IDHPIMTTTD FWTSHECLLL PYEQALTRED
 AtDHS3   DPQRMIRAYC QSAATLNLLR AFATGGYAAM QRVTQWNLDF TERSEQGDRY RELANRVDEA LGFMHAAGLT LDHPIMQTTD FWTSHECLLL PYEQSLTRLD
 AtDHS1   DPNRMIRAYT QSAATLNLLR AFATGGYAAI QRVTQWNLDF VEQSEQADRY QELANRVDEA LGFMSACGLG TDHPLMTTTD FYTSHECLLL PYEQSLTRLD
 AtDHS2   DPHRMVRAYT QSVATLNLLR AFATGGYAAM QRVSQWNLDF TQHSEQGDRY RELANRVDEA LGFMGAAGLT SAHPIMTTTE FWTSHECLLL PYEQALTRED

          301                                                                                                    400
OsDHS1b   STTGHFYDCS AHMLWVGERT RQLDGAHVEF LRGVANPLGI KVSDKMNPTE LVKLIEILNP SNKPGRITII TRMGAENMRV KLPHLIRAVR HAGQIVTWIT
BdDHS1b   STSGLFYDCS AHMLWVGERT RQLDGAHVEF LRGVANPLGI KVSDKMNPAD LVKLIDILNP TNKPGRITII TRMGAENMRV KLPHLIRAVR HAGQIVTWIT
PlDHS1b   STSGLFYDCS AHMLWVGERT RQLDGAHVEF LRGIANPLGI KVSDKMNPTE LVRLIEILNP SNKPGRITII TRMGAENMRV KLPHLIRAVR RAGQIVTWVT
BdDHS1a   STSGLFYDCS AHMLWVGERT RQLDGAHVEF LRGIANPLGI KVSDKMNPAE LVKLIEILNP SNKPGRITII TRMGAENMRV KLPHLIRAVR NSGQIVTWIT
 AtDHS3   STSGLYYDCS AHMIWVGERT RQLDGAHVEF LRGVANPLGI KVSDKMDPKE LVKLIEILNA DNKPGRITII TRMGAENMRV KLPHLIREVR RAGQIVTWVS
 AtDHS1   STSGLYYDCS AHMVWCGERT RQLDGAHVEF LRGIANPLGI KVSNKMDPFE LVKLVEILNP NNKPGRITVI VRMGAENMRV KLPHLIRAVR RSGQIVTWVC
 AtDHS2   STSGLYYDCS AHMLWVGERT RQLDGAHVEF LRGIANPLGI KVSDKMVPSE LVKLIEILNP QNKPGRITVI VRMGAENMRV KLPNLIRAVR GAGQIVTWVS

          401                                                                                                    500
OsDHS1b   DPMHGNTIKA PCGLKTRPFD SILAEVRAFF DVHDQEGSHP GGVHLEMTGQ NVTECIGGSR TVTFDDLGDR YHTHCDPRLN ASQSLELSFI IAERLRRKRI
BdDHS1b   DPMHGNTIKA PCGLKTRPFD SILAEVRAFF DVHEQEGSHA GGVHLEMTGQ NVTECIGGSR TVTFDDLGDR YHTHCDPRLN ASQSLELSFI IAEKLRKRRI
PlDHS1b   DPMHGNTIKA PCGLKTRPFD SILAEVRAFF DVHEQEGSHP GGVHLEMTGQ NVTECIGGSR TVTFDDLSDR YHTHCDPRLN ASQSLELAFI IAERLRKRRI
BdDHS1a   DPMHGNTIKA PCGLKTRPFD SIMNEVRAFF DVHDQEGSHP GGIHLEMTGQ NVTECIGGSR TVTFDDLGDR YHTHCDPRLN ASQSLELAFI IAERLRKRRM
 AtDHS3   DPMHGNTIKA PCGLKTRPFD AILAEVRAFF DVHEQEGSHP GGIHLEMTGQ NVTECIGGSR TVTFDDLGSR YHTHCDPRLN ASQSLELSFI IAERLRKRRI
 AtDHS1   DPMHGNTIKA PCGLKTRAFD SILAEVRAFL DVHEQEGSHA GGIHLEMTGQ NVTECIGGSR TVTYDDLSSR YHTHCDPRLN ASQSLELAFI VAERLRKRRT
 AtDHS2   DPMHGNTIMA PGGLKTRSFD AIRAELRAFF DVHDQEGSFP GGVHLEMTGQ NVTECVGGSR TITYNDLSSR YHTHCDPRLN ASQSLELAFI IAERLRKRRL

          501        516
OsDHS1b   RSSKLNNMLP LPPFGV
BdDHS1b   RSSKLNSVLP LPTYGF
PlDHS1b   RSSGLNNVLP LPPLAF
BdDHS1a   QS-GLTNNLP LPPLAF
 AtDHS3   KSQKAFAL
 AtDHS1   GSQRVS
 AtDHS2   GSGNLPSSIG V
```

**Supplemental Figure S9**. **Multiple sequence alignment of deregulated BdDHS1b and OsDHS1b, compared to Arabidopsis DHSs.** The *sota* domain region (marked in grey), which concentrates a high frequence of mutations that deregulate Arabidopsis DHSs (Yokoyama et al., 2022), is largely conserved in grass deregulated enzymes. Other residues affected by *sota* mutations deregulating Arabidopsis DHSs outside the *sota* region are marked in yellow, and are also largely conserved.

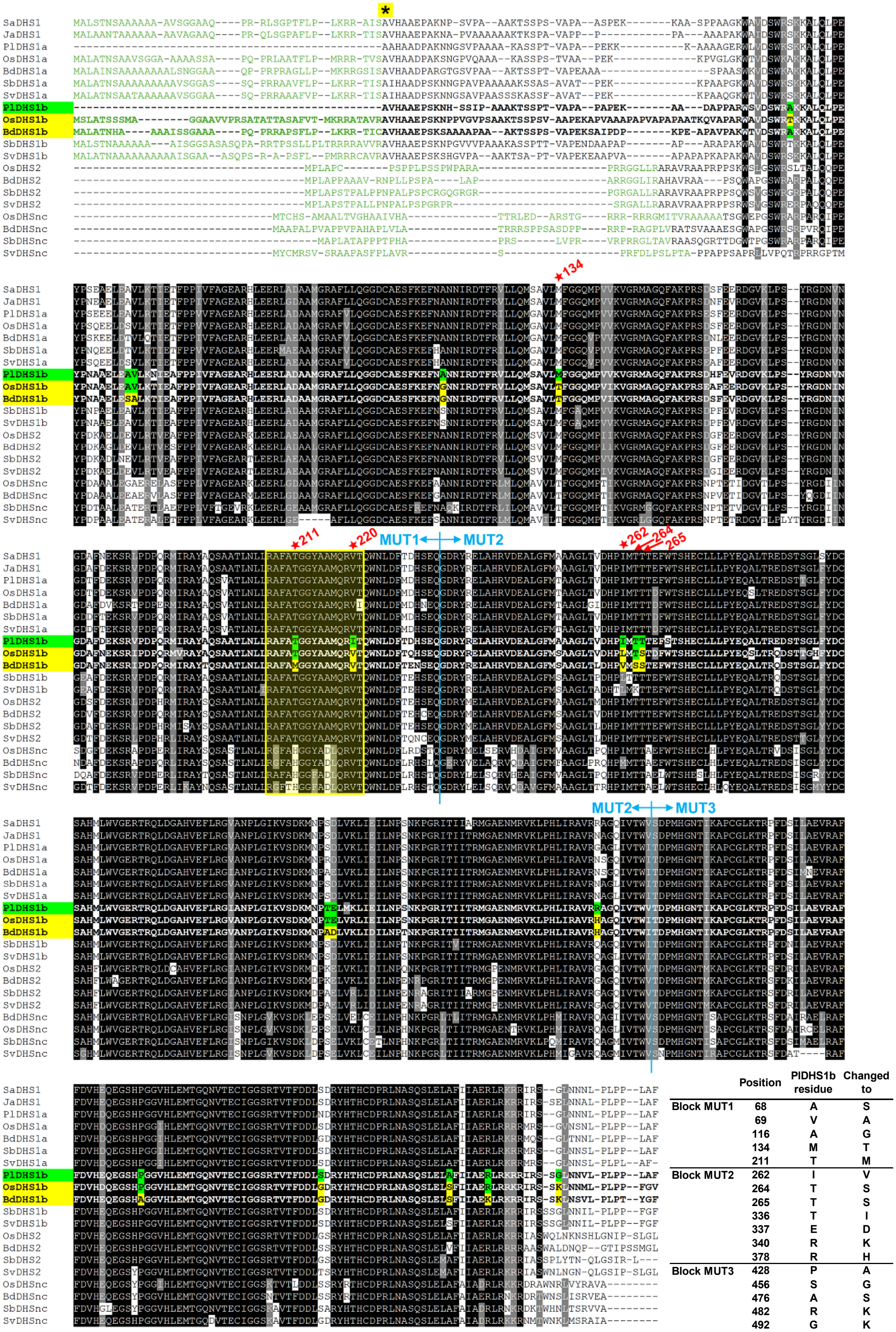


| | Position | PlDHS1b residue | Changed to |
|---|---|---|---|
| Block MUT1 | 68 | A | S |
| | 69 | V | A |
| | 116 | A | G |
| | 134 | M | T |
| | 211 | T | M |
| Block MUT2 | 262 | I | V |
| | 264 | T | S |
| | 265 | T | S |
| | 336 | T | I |
| | 337 | E | D |
| | 340 | R | K |
| | 378 | R | H |
| Block MUT3 | 428 | P | A |
| | 456 | S | G |
| | 476 | A | S |
| | 482 | R | K |
| | 492 | G | K |

**Supplemental Figure S10 (Previous page)**. **Multiple sequence alignment of DHS proteins from graminids.** Key residues targeted for mutagenesis in *Pharus lappulaceous* DHS1b sequence (PlDHS1b) are highlighted in green. The equivalent residues on the de-regulated DHSs BdDHS1b and OsDHS1b are marked in yellow. Vetical blue lines mark the transition between the groups of residues independently tested in blocks in $PlDHS1^{MUT1}$, $PlDHS1^{MUT2}$ and $PlDHS1^{MUT3}$. Key residues in positions 134, 211, 220, 262, 264, and 265 are marked with red stars/arrows. Positions have been numbered based on PlDHS1b sequence without its predicted trasit peptide (marked as green letters at the begining of the alignment). The asterisk highlighted in yellow on the first line of the alignment marks position 2 within PlDHS1b sequence. Note that *Pharus lappulaceous PlDHS1a* and *PlDHS1b* were directly cloned without the predicted transit peptide region using the orthologs from the closely related species *Pharus latifolius for* primer design, therefore the exact sequence of these predicted transit peptides is unknown. The region shadowed in pale yellow between residues 207 and 221 corresponds to the *sota* domain, previously identified in a supressor screening of Arabidopsis (Yokoyama et al., 2022).

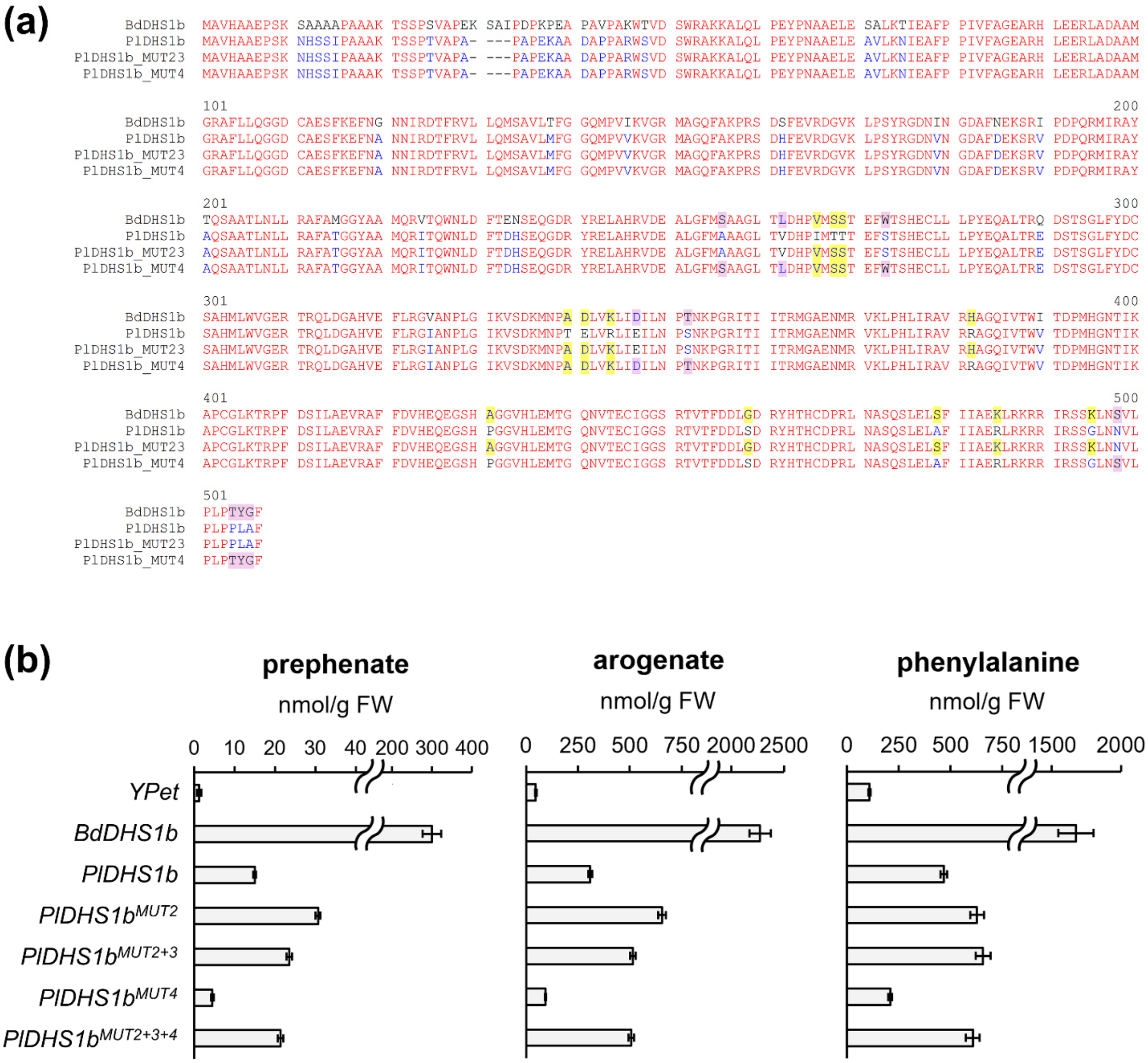


**Supplemental Figure S11. Transient overexpression of *PlDHS1b* with additional MUT4 mutations. (a)** Multiple sequence aligment comparing BdDHS1b, PlDHS1b, the double mutant MUT2+MUT3 (residues marked in yellow) constructed by assembling the mutant gBlocks MUT2 and MUT3, and the novel MUT4 (residues marked in purple). Note that residues at positions 262, 264, 265, 336, 337 and 340 of PlDHS1b sequence are redundant between MUT2+3 and MUT4. **(b)** Quantification of metabolites after transient expression of the newly generated PlDHS1b mutated proteins in *Nicotiana benthamiana*. Bars are the average of n = 5 replicates from independent plants, error bars = *SE*.

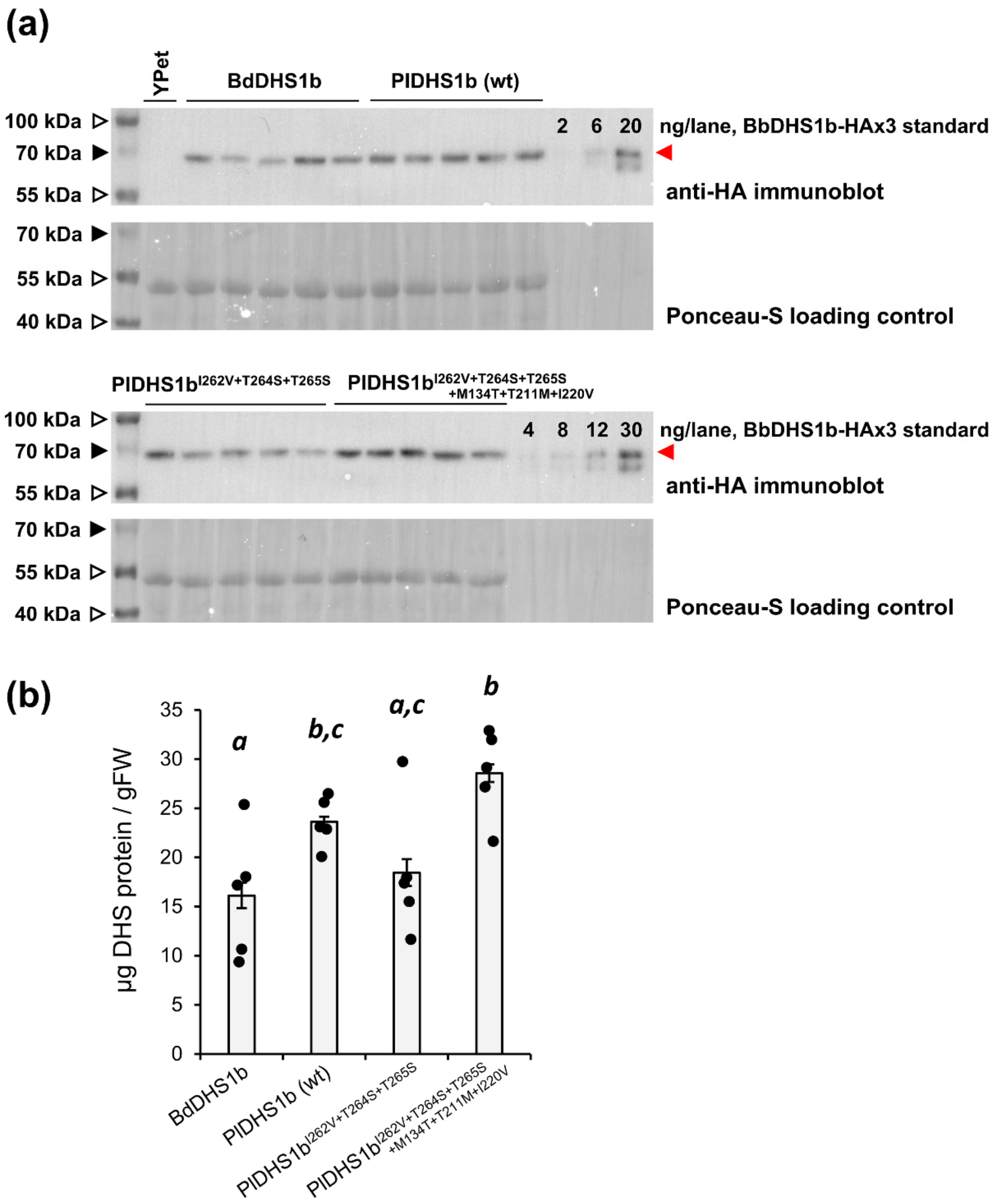


**Supplemental Figure S12**. **Immunoblot of *Pharus lappulaceous* DHS1b wild type and mutated proteins C-terminal 3xHA fusion proteins in *Nicotiana benthamiana*. (a)** Anti-HA immunoblot and on-membrane Ponceau S staining of total proteins extracted from plant simples. A standard curve was generated recombinant BdDHS1b-3xHA protein (mass in ng indicated above each lane; only the upper BdDHS1b-3xHA band was used for quantification). Each lane corresponds to an independent biological sample. Red arrows indicate the expected size of the mature protein without transit peptide. The images correspond to two SDS-PAGE gels that were transferred to a single membrane analyzed and analyzed in the same western blot experiment. **(b)** Quantification of DHS abundance according to the results shown in (a) and normalized by mass of plant tissue extracted (*n* = 5; error bars = *SE*).
